# Projection-Targeted Photopharmacology Reveals Distinct Anxiolytic Roles for Presynaptic mGluR2 in Prefrontal- and Insula-Amygdala Synapses

**DOI:** 10.1101/2024.01.15.575699

**Authors:** Hermany Munguba, Vanessa A. Gutzeit, Ipsit Srivastava, Melanie Kristt, Ashna Singh, Akshara Vijay, Anisul Arefin, Sonal Thukral, Johannes Broichhagen, Joseph M. Stujenske, Conor Liston, Joshua Levitz

## Abstract

Dissecting how membrane receptors regulate neural circuit function is critical for deciphering basic principles of neuromodulation and mechanisms of therapeutic drug action. Classical pharmacological and genetic approaches are not well-equipped to untangle the roles of specific receptor populations, especially in long-range projections which coordinate communication between brain regions. Here we use viral tracing, electrophysiological, optogenetic, and photopharmacological approaches to determine how presynaptic metabotropic glutamate receptor 2 (mGluR2) activation in the basolateral amygdala (BLA) alters anxiety-related behavior. We find that mGluR2-expressing neurons from the ventromedial prefrontal cortex (vmPFC) and posterior insular cortex (pIC) preferentially target distinct cell types and subregions of the BLA to regulate different forms of avoidant behavior. Using projection-specific photopharmacological activation, we find that mGluR2-mediated presynaptic inhibition of vmPFC-BLA, but not pIC-BLA, connections can produce long-lasting decreases in spatial avoidance. In contrast, presynaptic inhibition of pIC-BLA connections decreased social avoidance, novelty-induced hypophagia, and increased exploratory behavior without impairing working memory, establishing this projection as a novel target for the treatment of anxiety disorders. Overall, this work reveals new aspects of BLA neuromodulation with therapeutic implications while establishing a powerful approach for optical mapping of drug action via photopharmacology.

**Highlights:**

- Basolateral amygdala is a key site for anxiolytic action of mGluR2 agonism
- BLA receives dense Grm2^+^ inputs from ventromedial prefrontal cortex and posterior insular cortex
- Photoactivation of mGluR2 has distinct anxiolytic effects in vmPFC-BLA and pIC-BLA synapses
- Grm2^+^ vmPFC and pIC projections target medial and lateral BLA subregions, respectively

## Introduction

G protein-coupled receptors (GPCRs) serve as the largest class of drug targets for neurological and psychiatric disorders,^1,2^ motivating a deep understanding of their mechanisms of action. At the cellular level GPCRs operate via initiation of intracellular signaling cascades^3,4^ which can powerfully modulate neuronal activity and synaptic strength. Despite great progress in deciphering the structural basis of GPCR-mediated drug action,^5,6^ it has been difficult to understand receptor action in the functional context of complex neural circuits which, ultimately, mediate brain function and behavior. Central brain hubs, such as the amygdala, send and receive projection to and from various brain regions, raising questions about how different GPCR subtypes exert their actions on specific circuit elements to drive both therapeutic efficacy and side effects.

Anxiety disorders, including general anxiety disorder, panic disorder, phobias, agoraphobia, and social anxiety disorder, represent related neuropsychiatric diseases that, while heterogenous in the triggers and nature of their symptoms, are characterized by excessive worry and maladaptive defensive behaviors.^7,8^ Current therapeutic approaches for these disorders are limited in efficacy and can produce undesirable side effects.^9,10^ An integrative understanding of the molecular, synaptic and circuit mechanisms regulating avoidant and defensive behavioral phenotypes is needed to develop improved treatments for anxiety disorders.

The amygdala is a critical mediator of fear and anxiety, and efforts to develop treatments for anxiety disorders have revolved around normalizing its activity. The basolateral amygdala (BLA), a cortex-like amygdala sub-structure, serves as a central gate to relay sensory and cognitive information to other amygdala nuclei and limbic structures involved in emotional processing.^11^ Notably, direct optogenetic stimulation of BLA pyramidal neurons drives avoidance behavior.^12,13^ The BLA receives myriad cortical inputs, but the ways in which different cortical regions contribute to top-down control of the BLA is not well understood. For example, the ventromedial prefrontal cortex (PFC), a region highly interconnected with the BLA, has been reported to have both anxiogenic^14–16^ and anxiolytic^17,18^ roles. The insular cortex (IC) is also interconnected with the amygdala^19,20^ and has emerged as a bridge between sensation, interoception, and emotional processing.^21–26^ Notably, in humans, enhanced IC-BLA resting-state functional connectivity is observed in states of anxiety,^27^ suggesting that the IC may be a driver of the enhanced BLA activity observed in anxiety and fear disorders.^28^

The metabotropic glutamate receptors (mGluRs) form an eight-member GPCR family that are expressed throughout the nervous system where they respond to patterns of extracellular glutamate to tune synapse function over time through various forms of plasticity.^29,30^ Despite a wealth of preclinical and clinical data supporting the potential of mGluRs as therapeutic targets for the treatment of a wide range of neurological and psychiatric diseases,^31^ a circuit- and synapse-level view of mGluR-mediated control of behavioral processes is limited. mGluR2 has been highly implicated in the pathophysiology and treatment of anxiety disorders^32,33^ with preclinical rodent studies showing anti-avoidance effects of non-specific mGluR2/3 agonists^34^ without the sleep-inducing or addictive effects of benzodiazepines.^35^ Human trials with mGluR2/3 agonists or mGluR2-specific positive allosteric modulators have shown efficacy in general anxiety disorder^36,37^ and, most recently, in panic disorder^38^. Despite these promising results, mechanistic studies of mGluR2-mediated regulation of anxiety have been limited by the lack of precise means of manipulating specific receptor subtypes in genetically defined cells or projections with spatiotemporal precision.

While evidence exists that mGluR2 is present in presynaptic terminals in the BLA^39,40^ and electrophysiological studies have shown that mGluR2/3 agonists can drive long-lasting presynaptic inhibition in the BLA,^41–43^ the specific circuits regulated by mGluR2 remain unknown. Prior studies have also been limited by drugs that are unable to distinguish pre-synaptic versus post-synaptic or glial receptors, cannot be rapidly applied and removed, and typically target both mGluR2 and mGluR3. Here we demonstrate that local mGluR2/3 agonism in the BLA mediates an anxiolytic-like effect on anxiety-related behavior, and we subsequently dissect the role of mGluR2 signaling at specific synaptic inputs to the BLA. Using viral tracing, we demonstrate that mGluR2-expressing projections to the BLA arise predominantly from the ventromedial PFC and posterior IC. Using optogenetics-assisted electrophysiology and photopharmacology, we uncover the distinct anatomical, synaptic, and electrophysiological properties of these projection classes and reveal complementary but distinct roles in the regulation of anxiety-associated behavior. Together this work represents a novel strategy for delineating the underlying mechanisms of GPCR-targeting drug action within a complex neural circuit.

## Results

### mGluR2/3 activation in the BLA recapitulates behavioral effects of systemic activation

As a first assessment of the effects of mGluR2 on anxiety-associated behaviors, we intraperitoneally (i.p.) injected either the mGluR2/3-selective antagonist, LY341495 (LY34), or the mGluR2/3-specific agonist, LY379268 (LY37), and measured spatial avoidance in adult male mice using the open field test (OFT) and elevated plus maze (EPM). In both assays, LY34 increased and LY37 decreased avoidance behavior (**Fig. 1A, B**). In the OFT, LY37 increased the time spent in the center area of the chamber with the size of the effect increasing over the course of the 20 min experiment (**Fig. S1A**). LY34 had the opposite effect on time spent in the center (**Fig. 1A; S1A**) and decreased both the overall distance travelled (**Fig. S1B, C**) and the number of entries into the center area (**Fig. S1D**). In the EPM, LY37 increased while LY34 decreased the time spent in the open arm. Neither drug altered the overall locomotion (**Fig. S1E**), but LY37 increased the number of open arm entries (**Fig. S1F**) while LY34 strongly decreased the number of head dips (**Fig. S1G**). Importantly, LY37 had similar effects in EPM when the assay was performed with female mice (**Fig. S1H-K**).

**Figure 1.**
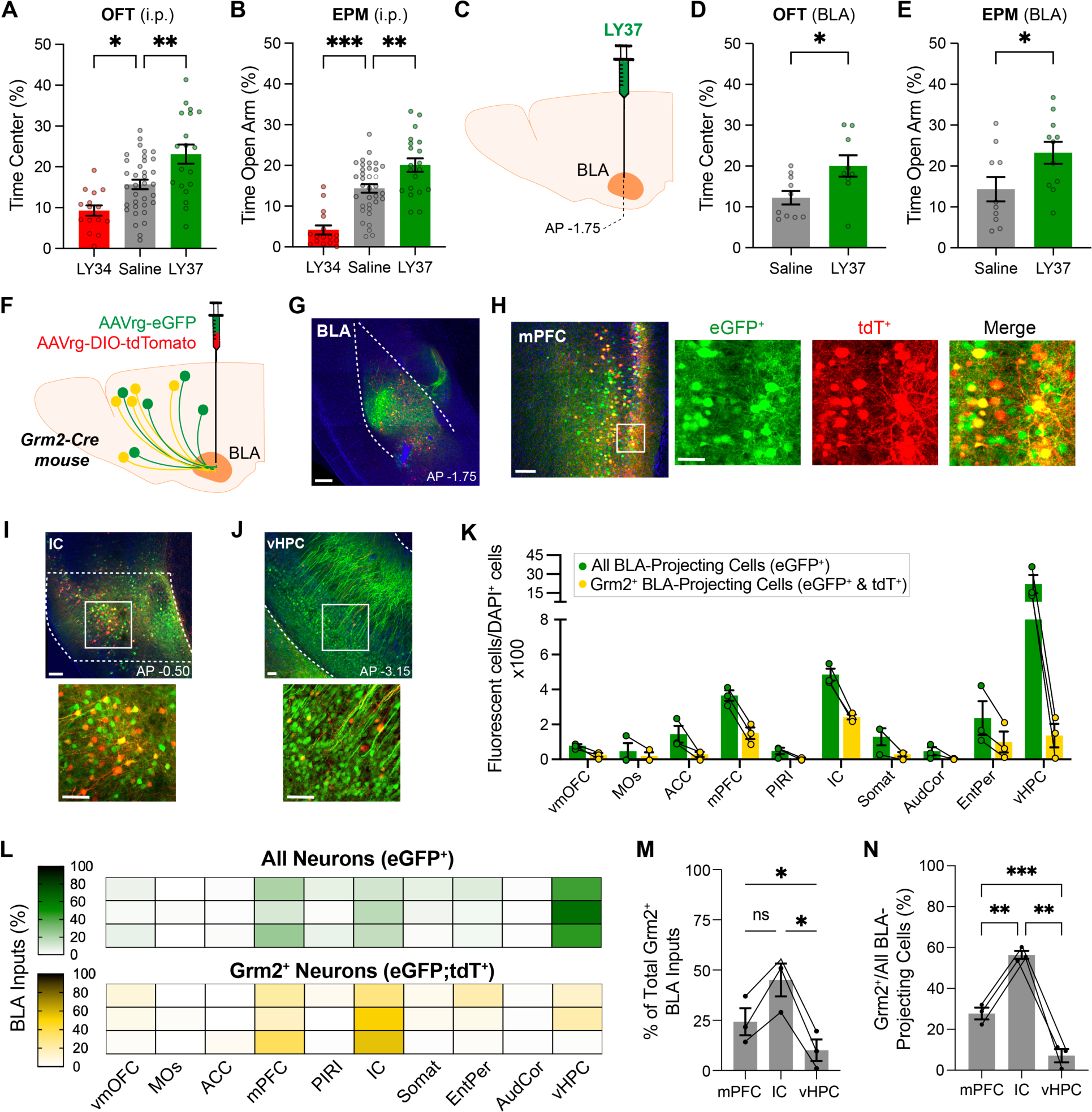
Anxiolytic effects of mGluR2/3 agonism in the BLA and mapping Grm2^+^ cortico-amygdalar projections. **(A-B)** I.P. injection of mGluR2/3 antagonist LY341495 (3 mg/kg) or agonist LY379268 (3 mg/kg) decreased and increased, respectively, spatial avoidance in the OFT (A) and EPM (B). **(C-E)** Intra-BLA infusion (C) of LY37 decreased spatial avoidance in the OFT (D) and EPM (E) compared to saline controls. **(F)** Dual injection of AAVrg-eGFP and AAVrg-DIO-tdTomato into the BLA of Grm2-Cre mice were performed to define BLA-projecting cells throughout the brain. **(G-J)** Representative images of dual retrograde virus expression at the BLA injection cite (G), and the medial prefrontal cortex (H), insular cortex (I) and ventral hippocampus (J). Zoomed in images show eGFP^+^ (green; total BLA-projecting cell population) and tdT^+^ (red; Grm2+ BLA-projecting cell population). **(K)** Summary graph of the density of total (green) and Grm2+ (yellow) BLA projecting cells across cortical regions normalized to total DAPI+ cells including the ventral medial orbitofrontal cortex (vmOFC), primary motor cortex (MOs), anterior cingulate cortex (ACC), mPFC, IC, somatosensory cortex (Somat), auditory cortex (AudCor), entorhinal and perirhinal cortex (EntPer), and ventral hippocampus (vHPC). numbers normalized to DAPI^+^ cells per region. (**L**) Heat maps showing fraction of BLA-targeting cells across cortical region separated by mouse (rows). Analysis of all cells (top) show vHPC as the most abundant input area, while analysis of Grm2^+^ cells (bottom) show the mPFC and IC as the most abundant input areas. **(M)** Summary bar graph of the perecentage of the total Grm2^+^ cortical BLA inputs from (L) graphed for mPFC, IC, and vHPC. **(N)** Comparison of the percentage of BLA-projecting cells which are Grm2^+^ (yellow/green *100) in mPFC, IC, and vHPC. For (A), (B), (D), (E), (K), (M), (N): Points represent individual mice. For (A), (B), (M), (N) 1-way ANOVA was used and for (D), (E) unpaired t-test is used. All data shown as mean ± SEM; * P<0.05, ** P < 0.01, *** P < 0.001. Scale bars are 200 µm, except for the inset zoomed images where it is 20 µm. See also Figure S1.

We next asked if the avoidance-decreasing effects of LY37 may be mediated by drug action in the amygdala by directly infusing LY37 into the BLA immediately prior to either the OFT or EPM assays. In both cases, local LY37 infusion recapitulated the anti-avoidance effects of systemic injection of LY37 without locomotion effects (**Fig. 1C, D; Fig. S1L-R**), establishing the BLA as a primary target of mGluR2/3-mediated regulation of anxiety-related behavior.

### Grm2^+^ cortical inputs to the BLA are enriched in ventromedial PFC and posterior IC

Given the key role of cortical inputs in controlling BLA activity and that mGluR2 is a largely presynaptic receptor that modulates cortico-amygdalar transmission,^39,41–44^ we asked which BLA-targeting projections contain mGluR2 as these are likely candidates for mediating the anxiolytic effect of LY37. We took advantage of Grm2-Cre mice which we recently established as a reliable reporter of mGluR2 expression^45^ that enables targeted expression in the subset of neurons which natively express mGluR2. Grm2-Cre mice were unilaterally injected into the BLA with a mixture of a Cre-dependent retrograde adeno-associated virus (rgAAV)-tdTomato (tdT), to label BLA-projecting Grm2^+^ cells, and a rgAAV-CAG-eGFP, to target all BLA-projecting cells. Following six weeks of expression, this revealed a brain-wide map of regions that target the BLA and labeled subsets of BLA-projecting cells that are also mGluR2-expressing (**Fig. 1F, G**). Quantification across all cortical regions revealed expression of BLA-projecting cells across many regions (**Fig. 1K-N; Fig. S1S-V**), with the highest density of BLA-projecting cells being observed in the medial PFC (mPFC), IC, and ventral hippocampus (vHPC) (**Fig. 1H-J**). However, only the mPFC and IC showed a large proportion of Grm2^+^ (tdT^+^) BLA projecting cells (**Fig. 1H-J**), consistent with prior work showing weak expression of mGluR2 in hippocampus.^46^ Grm2^+^ BLA-projecting cells were also seen in subcortical regions, including contralateral BLA (**Fig. S1W**).

Based on these results, we decided to focus our analyses on mPFC-BLA and IC-BLA projections as these represent the vast majority of Grm2^+^ BLA-projecting cortical neurons. Closer analysis of rAAV-driven fluorescent protein expression in mPFC revealed that Grm2^+^ BLA-projecting cells were enriched in vmPFC with progressively decreased density in dorsomedial PFC and anterior cingulate cortex (ACC) (**Fig. 2A-C; Fig. S2A**). Grm2^+^ BLA-projecting cells were enriched in layer 2/3 compared to deeper layers (**Fig. 2D**). While ∼50% of BLA-projecting cells in layer 2/3 were Grm2^+^, less than 25% were Grm2^+^ in layer 5 and layer 6 (**Fig. 2E**). A small number of green and yellow cell bodies were observed in the contralateral ventromedial PFC (vmPFC) (**Fig. S2B**), suggesting that vmPFC-BLA projections are primarily ipsilateral.

**Figure 2.**
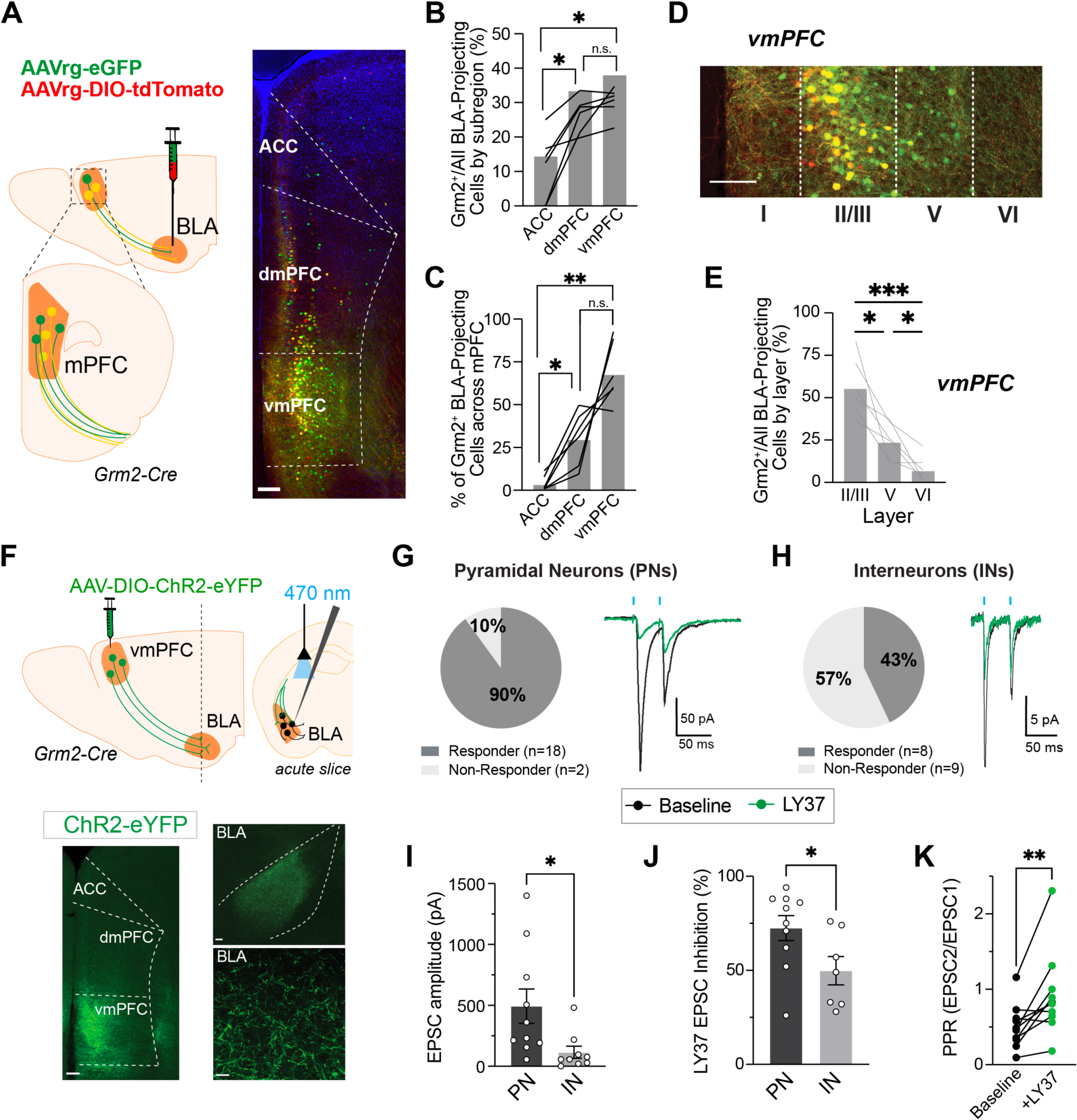
Grm2^+^ inputs to the BLA are enriched in layer 2/3 of the vmPFC and show mGluR2/3 agonist sensitivity. **(A-C)** Representative image (A) and analysis (B-C) showing BLA-projecting cells in mPFC subregions: anterior cingulate cortex (ACC), dorsal medial PFC (dmPFC), and ventral medial PFC (vmPFC). A higher percentage of BLA-projecting cells are Grm2^+^ in vmFPC compared to dmPFC and ACC (B) and vmPFC contains the majority of Grm2^+^ BLA-projecting cells across the mPFC (C). **(D-E)** Representative image and analysis showing that Grm2^+^ BLA-projecting cells are enriched in layer 2 and 3 of the vmPFC. A higher percentage of BLA-projecting cells is Grm2^+^ in layer 2/3 compared to deeper layers (E). **(F)** Schematic (top) and representative images (bottom) showing optogenetics-assisted analysis of Grm2^+^ vmPFC to BLA projections. **(G-H)** Pie charts (left) and representative images (right) showing connectivity between Grm2^+^ vmPFC projections and either pyramidal neurons (PNs; G) or interneurons (INs; H). Application of LY37 (green traces) decreases optical EPSC (oEPSC) amplitude. Blue lines indicate 1 ms 470 nm light pulses at 20 Hz to stimulate axonal ChR2 and elicit neurotransmitter release. **(I-K)** Bar graphs showing that larger oEPSCs are seen in PNs (I), the inhibitory effect of LY37 is larger in PNs (J), and LY37 produces an increase in paired pulse ratio (PPR) in PNs (K). Points represent individual mice (B, C, E) or individual cells (K, L). For (B), (C), (E) 1-way ANOVA was used; for (I), (J) unpaired t-test was used; for (K) paired t-test was used. All data shown as mean ± SEM; * P<0.05, ** P < 0.01, *** P < 0.001. Scale bars are 200 µm (A, left image in F) or 20 µm (D, right images in F). See also Figure S2.

To directly assess connectivity between Grm2^+^ vmPFC cells and BLA, we unilaterally injected a Cre-dependent AAV for ChR2 in the vmPFC of Grm2-Cre mice and recorded optically evoked EPSCs (oEPSCs) in acute slices of the BLA (**Fig. 2F**). 90% of pyramidal neurons (PNs) and <50% of interneurons (INs), as determined by cell morphology, showed oEPSCs (**Fig. 2G, H**), with substantially larger amplitudes in PNs (**Fig. 2I**). Inputs to both PNs and INs were highly sensitive to LY37 application (**Fig. 2G, H, J; Fig. S2C-E**), though the effect of LY37 was larger in PNs (**Fig. 2J**), which showed a clear increase in paired pulse ratio (**Fig. 2K**), indicative of a presynaptic mechanism of inhibition. Interestingly, the latency from stimulation to oEPSC onset was larger for PNs than INs (**Fig. S2F**). Similar results were seen in the presence of tetrodotoxin (TTX) (**Fig. S2G-K**), confirming the monosynaptic nature of Grm2^+^ vmPFC-BLA projections.

Analysis of rAAV-driven fluorescent protein expression in the IC revealed a modest enrichment of Grm2^+^ projections in the posterior IC (pIC) relative to medial IC (mIC) and anterior IC (aIC) (**Fig. 3A-C**). Within the pIC, Grm2^+^ projections were enriched in the agranular subregion (**Fig. 3D, E; Fig. S3A**). Grm2^+^ BLA-projecting cells in the pIC showed largely the same laminar distribution as the total population of BLA-projecting cells, which were distributed across layers (**Fig. S3B, C**). Unlike the vmPFC projections, a substantial population of cell bodies were observed in the contralateral pIC (**Fig. S3D**), suggesting that pIC-BLA projections are intra- and inter-hemispheric. Using the same ChR2-based functional mapping approach as used for the vmPFC, we found that Grm2^+^ pIC projections targeted both PNs and INs but with higher connectivity and oEPSC amplitude for PNs (**Fig. 3F-I**). LY37 also produced a strong presynaptic inhibition of pIC-BLA oEPSCs (**Fig. 3G, H, J, K; Fig. S3E-G**), though with a comparable effect in PNs and INs (**Fig. 3J**). Unlike vmPFC-BLA projections, a similar oEPSC latency was seen in PNs and INs (**Fig. S3H**).

**Figure 3.**
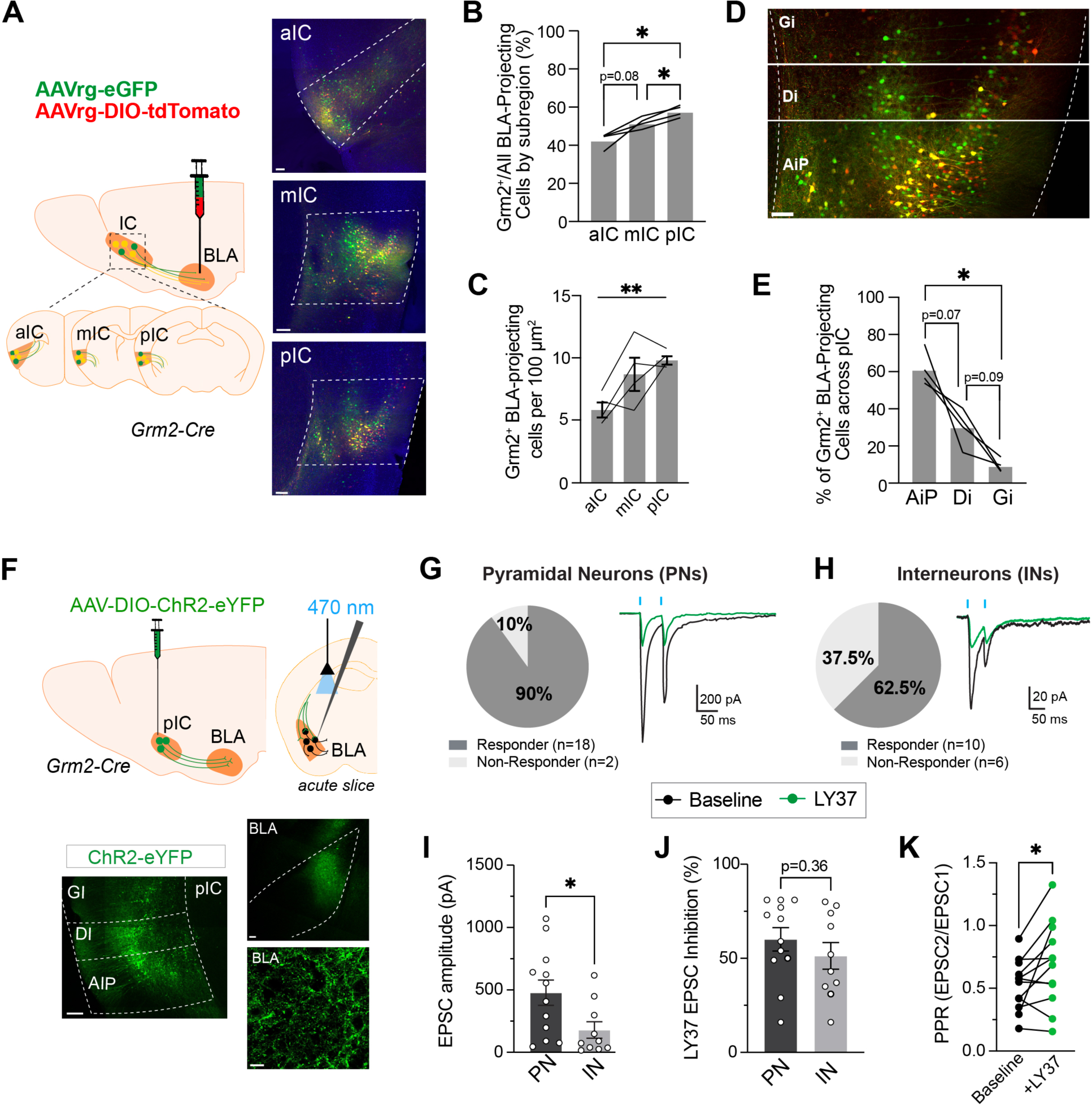
Grm2^+^ inputs to the BLA are enriched in the agranular subregion of the pIC and show mGluR2/3 agonist sensitivity. **(A-C)** Representative images (A) and analysis (B-C) showing BLA-projecting cells in IC subregions: anterior (aIC), medial (mIC), and posterior (pIC). A higher percentage of BLA-projecting cells are Grm2^+^ in pIC compared to aIC and mIC (B) and pIC contains a higher density of Grm2^+^ BLA-projecting cells (C). **(D-E)** Representative image (D) and analysis (E) showing that Grm2^+^ BLA-projecting cells are enriched in the agranular (AiP) subregion of the pIC compared to the dysgranular (Di) and granular (Gi) subregions. The largest population of Grm2^+^ BLA-projecting cells comes from AiP (E). **(F)** Schematic (top) and representative images (bottom) showing optogenetics-assisted analysis of Grm2^+^ pIC to BLA projections. **(G-H)** Pie charts (left) and representative images (right) showing connectivity between Grm2^+^ pIC projections and either pyramidal neurons (PNs; G) or interneurons (INs; H). Application of LY37 (green traces) decreases oEPSC amplitude. Blue lines indicate 1 ms 470 nm light pulses at 20 Hz to stimulate axonal ChR2 and elicit neurotransmitter release. **(I-K)** Bar graphs showing that larger oEPSCs are seen in PNs (I), the inhibitory effect of LY37 is comparable in PNs and INs (J), and LY37 produces an increase in paired pulse ratio (PPR) in PNs (K). Points represent individual mice (B, C, E) or individual cells (K, L). For (B), (C), (E) 1-way ANOVA was used; for (I), (J) unpaired t-test was used; for (K) paired t-test was used. All data shown as mean ± SEM; * P<0.05, ** P < 0.01. Scale bars are 200 µm (A, left image in F) or 20 µm (D, right images in F). See also Figure S3.

### Photo-activation of mGluR2 in Grm2^+^ vmPFC-BLA synapses decreases spatial avoidance

Based on the density of Grm2^+^ projections from the vmPFC and pIC to the BLA (**Fig. 2, 3**), the sensitivity of these synapses to mGluR2 activation (**Fig. 2, 3**), and the behavioral effect of intra-BLA LY37 infusion (**Fig. 1**), we hypothesized that presynaptic mGluR2 activation in either, or both, of these inputs drives the acute anxiolytic effects of mGluR2/3 agonists. To test this, we turned to a tethered photopharmacology approach which enables rapid optical control of specific receptors with genetic targeting *in vivo.*^45,47^ In brief, genetically encoded SNAP-tagged mGluR2 (SNAP-mGluR2) is expressed on the surface of cells where it can be covalently labeled by a “photoswitchable, orthogonal, remotely-tethered ligand” (PORTL) termed BGAG_12,400_ which allows photoactivation with 385 nm light that can be reversed by 525 nm light (**Fig. 4A**).^48^ Consistent with our prior work showing that SNAP-mGluR2 efficiently traffics and functions in presynaptic terminals,^45,49^ we expressed SNAP-mGluR2 via a Cre-dependent AAV in either vmPFC or pIC and observed clear expression both in cell bodies in the cortex and at terminals in the BLA via immunohistochemical labeling (**Fig. 4B, C; Fig. S4A-C**).

**Figure 4.**
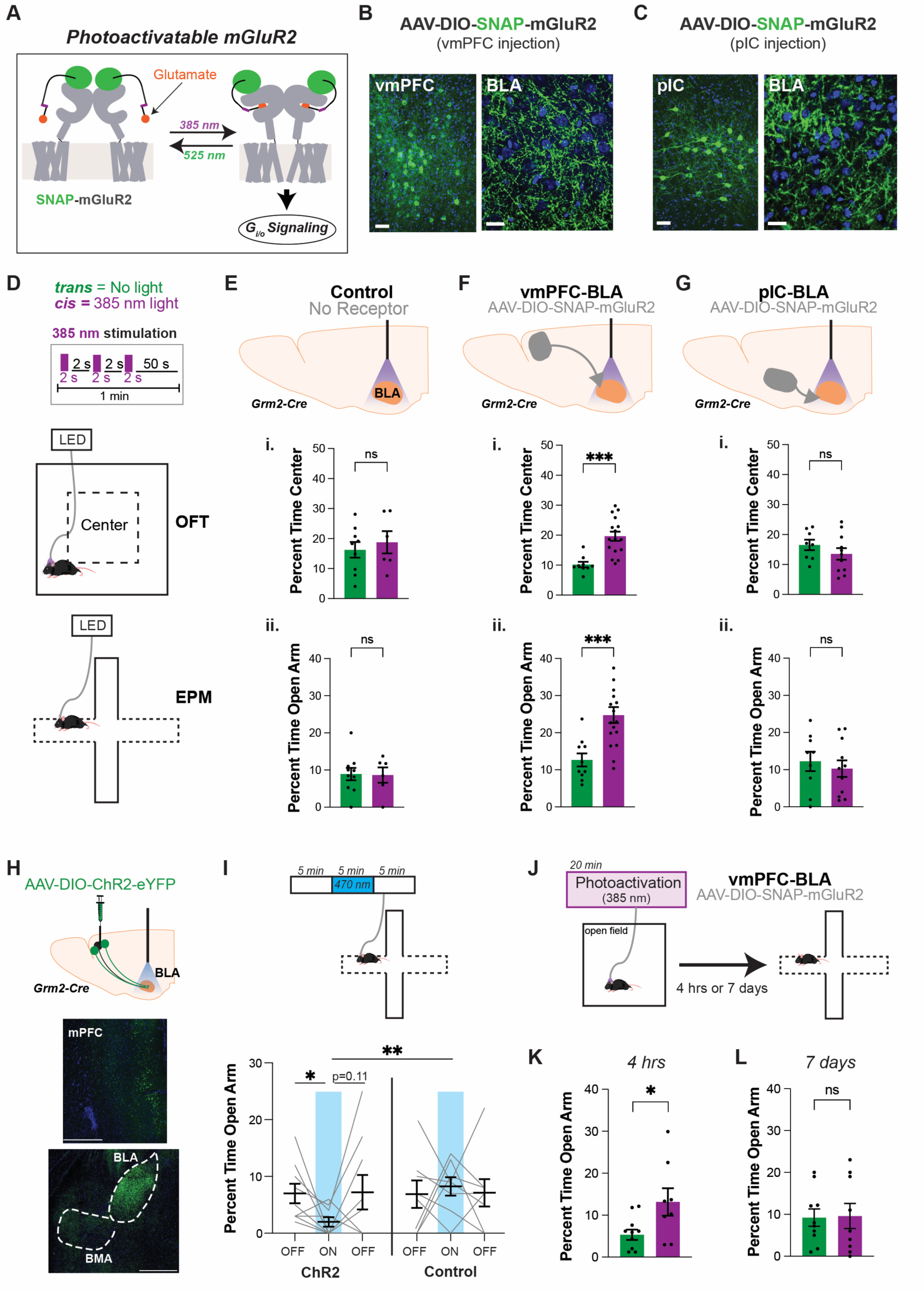
Presynaptic mGluR2 photoactivation in vmPFC-BLA synapses decreases spatial avoidance. **(A)** Schematic of photopharmacological mGluR2 photoactivation. N-terminally SNAP-tagged mGluR2 is expressed and covalently labeled with BGAG_12,400_, a photoswitchable, orthogonal, remotely-tethered ligand (PORTL). In its relaxed *trans* state BGAG_12, 400_ is inactive, but 385 nm illumination initiates conversion to the active *cis* state which drives agonism of mGluR2. Photoactivation is stable in the dark after 385 nm illumination but is reversed by visible light (>470 nm) illumination. **(B-C)** Representative images showing immunistochemical staining of an N-terminal HA-tag on SNAP-mGluR2, confirming robust expression (green) in cell bodies in the cortex (left) and processes in the BLA (right) in Grm2-Cre mice. DAPI staining is shown in blue. **(D)** Schematic showing *in vivo* mGluR2 photoactivation protocols for OFT and EPM experiments. **(E-G)** SNAP-mGluR2 photoactivation with 385 nm illumination (magenta) elicits a decrease in spatial avoidance compared to the no photoactivation condition (green) for vmPFC-BLA projection mice (F), but not for control (E) or pIC-BLA projection mice (G) in OFT (i) and EPM (ii). **(H)** Schematic and representative image showing ChR2-eYFP expression in Grm2^+^ vmPFC cell bodies and BLA terminals, but not in Basomedial Amygdala (BMA). **(I)** Schematic of EPM experiment (top) and summary graph (bottom) showing anxiogenic effect of vmPFC-BLA optogenetic stimulation (blue) in ChR2-expressing, but not control, mice. **(J)** Schematic showing behavioral plasticity experiment. SNAP-mGluR2 photoactivation occurs in the OFT for 20 min and EPM is measured 4 hrs or 7 days later. **(K-L)** Prior SNAP-mGluR2 photoactivation produces an anxiolytic effect in the EPM 4hrs (K), but not 7 days (L), later. Note that the control group in the 4 hr experiment showed a lower level of exploration (∼5% open arm time) compared to the 7-day experiment likely due to prior exposure to the anxiogenic open field 4 hours before the EPM. Points and lines represent individual mice. For (E-G), (K-L) unpaired t-test was used; For (I), 2-way ANOVA was used. All data shown as mean ± SEM; * P<0.05, **P<0.01, *** P < 0.001. Scale bars are 20 µm. See also Figure S4.

We then asked if *in vivo* photo-activation of presynaptic mGluR2 in either projection can recapitulate the behavioral effect of LY37. We prepared three groups of mice with expression of either SNAP-mGluR2 in the vmPFC or pIC or no receptor expression (**Fig. 4D-G**). All three groups received optic fiber/cannula implants targeting the BLA which were used to deliver BGAG_12,400_ ∼16 hr before the experiments and light during the experiment (**Fig. 4D; Fig. S4A, D**). Within each group, half of the mice received no light and the other half received 385 nm light to photo-activate mGluR2 during either OFT or EPM. All groups showed similar baseline values of center (OFT) or open arm (EPM) exploration and in both the control group and pIC group 385 nm light showed no effect (**Fig. 4E, G**). In contrast, 385 nm illumination led to an increase in center and open arm time for the vmPFC group (**Fig. 4F**), revealing that presynaptic mGluR2 photoactivation specifically at vmPFC-BLA synapses can drive a rapid decrease in spatial avoidance. mGluR2 photoactivation had no effect on locomotion in either group, but mGluR2 photoactivation in vmPFC projections did increase center entries in the OFT (**Fig. S4E**) and head dips and open arm entries in the EPM (**Fig. S4F**). Consistent with a net inhibitory effect, a decrease in BLA c-Fos immunostaining was observed after mGluR2 photoactivation in vmPFC projections (**Fig. S4G, H**).

Prior studies of vmPFC-amygdala projections have observed an anxiolytic or no effect of stimulation,^17,50,51^ seemingly at odds with our results showing that presynaptic inhibition of vmPFC-BLA projections is anxiolytic. One potential explanation is that, whereas prior studies stimulated the axons of a broad population of vmPFC neurons using a CaMKII promoter, we have targeted the Grm2^+^ subset of vmPFC-BLA axons. To test this, we bilaterally injected a Cre-dependent AAV for ChR2-eYFP in the vmPFC of Grm2-Cre mice and implanted optical fibers above the BLA where dense projections were observed (**Fig. 4H**). Compared to controls that received a Cre-dependent AAV for eGFP, optical stimulation in ChR2 group during the EPM task reversibly decreased the percentage of time in the open arm (**Fig. 4I**), the number of open arm entries (**Fig. S4I**), and the number of head dips (**Fig. S4J**). This result indicates that activity of Grm2^+^ vmPFC-BLA projections is indeed acutely anxiogenic, providing an explanation for the anxiolytic effect of presynaptic inhibition by mGluR2.

We then asked about the timescale of the behavioral effects of mGluR2-mediated inhibition of vmPFC-BLA synapses. Consistent with prior studies using electrical stimulation, LY37 application was able to drive a long-term depression (LTD) of vmPFC-BLA synapses that lasted for at least 30 min after drug removal (**Fig. S4K**). Motivated by this, we asked if SNAP-mGluR2 photo-activation in vmPFC-BLA synapses could lead to decreases in spatial avoidance that persist after receptor de-activation. To test this idea, we photo-activated SNAP-mGluR2 in vmPFC-BLA synapses for 20 min while mice freely explored an open field (**Fig. 4J**). At the end of 20 min, we deactivated SNAP-mGluR2 with 525 nm light and then performed EPM measurements either 4 hr or 7 days later. Strikingly, 4 hrs after SNAP-mGluR2 photoactivation mice showed increased open arm time, open arm entries, and head dips compared to mice that received BGAG_12,400_ injection and optic fiber attachment, but no photo-activation (**Fig. 4J,K; Fig. S4L**). This result indicates that long-lasting synaptic and circuit changes can occur after presynaptic mGluR2 activation which can extend behavioral control beyond the initial period of acute presynaptic inhibition. In contrast, 7 days following photo-activation no effect was seen (**Fig. 4J,L; Fig. S4M**).

### Distinct behavioral roles of vmPFC-BLA versus pIC-BLA projections

Given the dense projections from the pIC to the BLA (**Fig. 3**) and prior work implicating the pIC in negative valence processing,^19^ we hypothesized that Grm2^+^ pIC-BLA projections may play a larger role in forms of anxiety-related behaviors beyond spatial avoidance. We thus designed a behavioral battery to extend our photopharmacological analysis which included temporally spaced assessments (to minimize confounds) of a variety of relevant behaviors including OFT, EPM, a social interaction test (SIT), as a measure of social avoidance, novelty suppressed feeding (NSF), as a measure of hypophagia which incorporates prior food deprivation, fox urine avoidance test (FUAT), as a measure of exploratory behavior incorporating aversive olfactory input, the Y-maze test (YMT), as a measure of working memory, and the tail suspension test (TST), as a depression-related measure of active coping (**Fig. 5B**).

**Figure 5.**
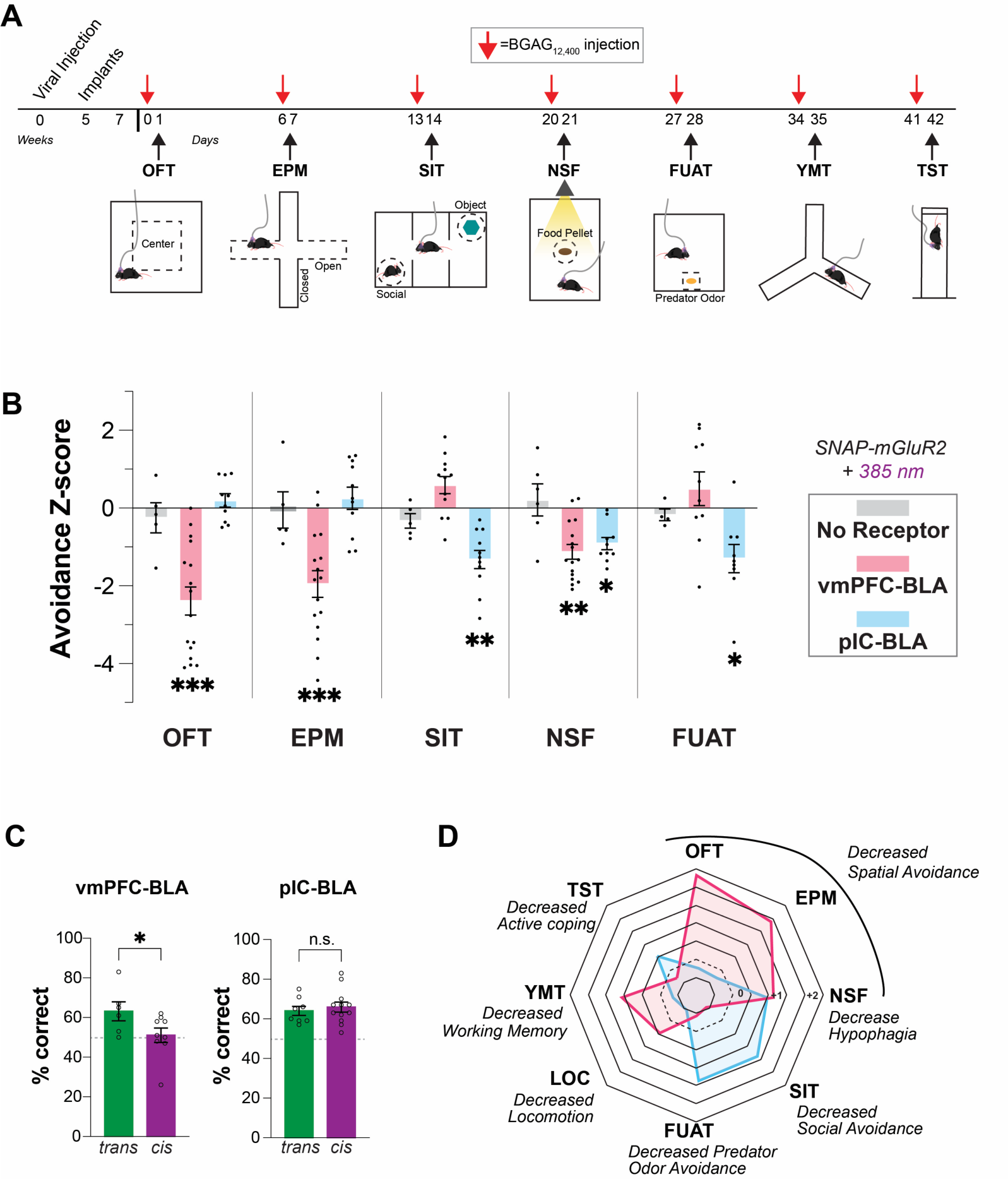
Distinct anxiolytic effects of SNAP-mGluR2 photo-activation in vmPFC-BLA versus pIC-BLA terminals reveals unique roles for each projection. **(A)** Timeline of avoidance behavior battery showing BGAG12,400 injection 1 day prior to each behavioral assay for each group (no SNAP-mGluR2, SNAP-mGluR2 in vmPFC-BLA projection, SNAP-mGluR2 in pIC-BLA projection). Each mouse either received no light for every assay or 385 nm for every assay. **(B)** Z-score analysis (see methods) showing effects of SNAP-mGluR2 photoactivation in each projection on avoidance behavior. **(C)** mGluR2 photoactivation in vmPFC-BLA (left), but not pIC-BLA (right), impairs working memory performance in the Y-maze test. **(D)** Radar plot summarizing the behavioral effects of SNAP-mGluR2 activation in either vmPFC-BLA (pink) or pIC-BLA (blue). Z-score values are used for each assay and locomotion data comes from OFT. Points represent individual mice. OFT = open field test, EPM = elevated plus maze, SIT = social interaction test, NSF = novelty suppressed feeding, FUAT = fox urine avoidance test, YMT = Y Maze Test, TST = tail suspension test. Statistics for corresponds to unpaired t-tests performed between light group and respective no-light control; Unpaired t-test was used in (C); All data shown as mean ± SEM; * P<0.05, ** P<0.01, *** P < 0.001. See also Figure S5.

Mice with SNAP-mGluR2 in vmPFC or PIC, as well as control mice that did not received SNAP-mGluR2, were tested in all behaviors with or without BLA-targeted photoactivation using a 7-day inter-behavior interval (**Fig. 5A, B**). Importantly, no light effects were seen in the control group across all assays (**Fig. S5A-E**). Unlike the EPM and OFT where only vmPFC-BLA photoactivation showed a clear effect (**Fig. 3**; **Fig. 5B**), SNAP-mGluR2 photoactivation in pIC-BLA synapses lead to increased social interaction (**Fig. 5B; Fig. S5A**). In contrast, SNAP-mGluR2 photoactivation in vmPFC-BLA synapses tended toward decreased social interaction (**Fig. S5A**), revealing a potential opposing effect of each projection. In the NSF test, SNAP-mGluR2 photoactivation strongly decreased the latency to feeding in the open field in either projection, while manipulation of the pIC-BLA pathway increased the latency to feeding both in the open field and in the home cage (**Fig. 5B; Fig. S5B**). The FUAT showed similar effects to the SIT, with increased and decreased exploratory behavior following SNAP-mGluR2 photoactivation in pIC-BLA and vmPFC-BLA, respectively (**Fig. 5B; Fig. S5C**). SNAP-mGluR2 photoactivation only in vmPFC-BLA synapses impaired working memory performance (**Fig. 5C**), while no effect was seen for either projection in the TST (**Fig. S5D**).

**Figure 5D** shows a summary of the behavioral battery results, highlighting the distinct effects of SNAP-mGluR2 photoactivation in either projection. Despite overlapping effects of mGluR2 photo-activation in vmPFC-BLA and pIC-BLA synapses in the NSF, they show a distinct profile of largely anxiolytic effects across the entire battery. vmPFC-BLA mGluR2 activation noticeably decreases spatial avoidance (OFT, EPM, NSF), and clearly impairs working memory (YMT), with minimal effects seen on social (SIT) or exploratory (FUAT) behaviors. In contrast, pIC-BLA mGluR2 activation generally increases exploration (SIT, FUAT), but only decreases spatial avoidance in the context of food seeking (NSF). Together these data suggest that distinct circuit organization of Grm2^+^ vmPFC-BLA and pIC-BLA projections enable them to drive distinct avoidance-related behaviors.

### Distinct organization of vmPFC-BLA and pIC-BLA circuits

Given the distinct behavioral effects of mGluR2 activation in vmPFC and pIC projections to BLA (**Fig. 5**), we hypothesized that the unique physiological roles of each projection may correspond to distinct connectivity patterns within the BLA. We first asked if Grm2^+^ vmPFC and Grm2^+^ pIC axons target the same or different cells within the BLA by expressing Cre-dependent spectrally separated opsins (ChR2-eYFP; Chrimson-tdTomato, see Methods) in either cortical region in the same Grm2-Cre mice (**Fig. 6A**). Axonal fluorescence in the BLA showed a spatial segregation of the two inputs with higher intensity for vmPFC projections in the medial parts of the BLA, sometimes described as anterior BLA (BLAa),^20,52^ and higher intensity for pIC projections in the lateral parts of the BLA (**Fig. 6B, C; Fig. S6A**), also described as posterior BLA (BLAp).^20,52^ In addition, rostral parts of the BLA showed vmPFC, but not pIC, innervation (**Fig. S6A**). Consistent with the high degree of connectivity observed in initial electrophysiology experiments using only one opsin (ChR2) (**Fig. 2, 3**), it was unsurprising that most cells (28/31 cells) showed oEPSCs following stimulation of both pathways (**Fig. 6D, E**). While, a wide range of relative amplitudes were seen, with some cells showing very strong inputs from both pathways and others showing much larger inputs from one pathway compared to the other (**Fig. 6E**), on average larger oEPSCs were observed for vmPFC inputs in the medial portions of the BLA (**Fig. 6G**), and larger oEPSCs for pIC inputs in lateral sections of the BLA (**Fig. 6F).** A non-significant trend was observed in terms of short-term plasticity, with a lower PPR observed for pIC inputs in the lateral BLA (**Fig. S6B**) and a lower PPR observed for vmPFC inputs in the medial BLA (**Fig. S6C**), suggesting that higher release probabilities contribute to the discrepancy in amplitudes between projection classes at each BLA subregion. Interestingly, a shorter latency of oEPSC onset was observed for pIC inputs to the lateral BLA compared to pIC inputs to the medial BLA or vmPFC inputs to either subregion (**Fig. S6D, E**). Together these data show that Grm2^+^ vmPFC and pIC projections target distinct but overlapping neuronal populations of the BLA, in line with their distinct roles in regulating avoidance behavior (**Fig. 5**).

**Figure 6.**
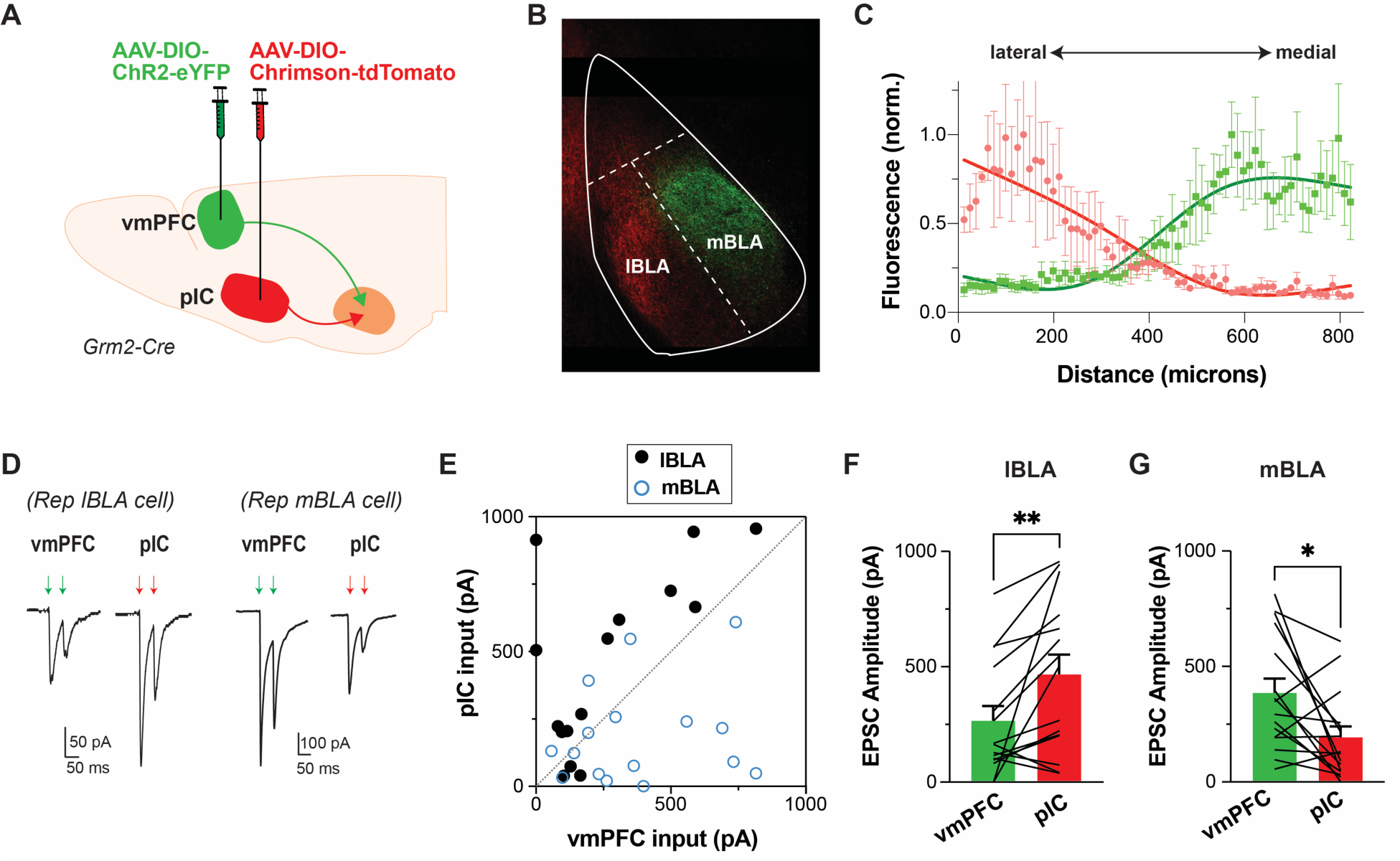
Overlapping but distinct BLA connectivity for Grm2^+^ vmPFC-BLA and pIC-BLA cells. **(A)** Schematic showing dual, two-color analysis of Grm2^+^ vmPFC and pIC inputs to the BLA. **(B-C)** Representative image (B) and quantification (C; average of 3 mice) showing that Grm2^+^ vmPFC inputs appear denser in medial BLA while Grm2^+^ pIC inputs appear denser in lateral BLA. **(D)** Representative traces showing relative size of synaptic input from vmPFC and pIC for a lateral and a medial BLA cell. **(E)** Summary scatter plot showing range of amplitude of oEPSCs from each input. Each point represents an individual cell in either the lateral or medial BLA. **(F-G)** Summary bar graph showing larger pIC-driven oEPSCs in lateral BLA (F) and larger vmPFC-driven oEPSCs in medial BLA (G). Paired t-test was used in (F, G); ** P<0.01. See also Figure S6.

## Discussion

Our work provides a novel approach to map the neural circuit basis of a receptor-targeting drug’s therapeutic action with high spatiotemporal and genetic precision. We first use targeted drug infusion to demonstrate that local mGluR2/3 agonism in the BLA is sufficient to induce an anxiolytic-like effect on spatial avoidance. Then, using cre-dependent viruses in Grm2-Cre mice, we pinpoint the receptor-expressing cells that can mediate the relevant behavioral effects of mGluR2 activation. We demonstrate that mGluR2/3 agonism induces presynaptic inhibition in Grm2^+^ projections from both the vmPFC or pIC. Using tethered photopharmacology to restrict receptor activation to Grm2^+^ vmPFC-BLA and pIC-BLA synapses for temporally precise periods, we show that mGluR2 activation on vmPFC or pIC presynaptic terminals induces anxiolytic-like effects on distinct sets of avoidance behaviors. Finally, we show that the amount of Grm2^+^ vmPFC and pIC input to the BLA exhibit virtually opposite mediolateral gradients, suggesting that at least part of the difference in the effect of mGluR2 activation in these two pathways is related to the downstream BLA circuits that they recruit.

This study represents the first application of tethered photopharmacology for projection-specific manipulation, and critically, it overcomes the lack of temporal precision afforded by genetic knockout or knock-in and the lack of spatial or genetic precision associated with pharmacology. Our work, thus, provides a template to manipulate the wild-type, full-length receptor of interest, which can be applied to map drug action with high spatiotemporal precision, rather than to merely manipulate the activity of a cell as is typical with traditional optogenetic or chemogenetic approaches. This approach holds promise for probing the circuit mechanisms mediating drug action for a range of other neuromodulatory GPCRs for which tethered photopharmacological tools^53–55^ and Cre-dependent reporter mice^56–60^ have been developed. One limitation of this approach is that it requires overexpression of a functional GPCR construct, but we demonstrate that this limitation can be overcome by carefully designed controls, as we did not find any influence of SNAP-mGluR2 expression in the absence of photoactivation. Approaches for pharmacological or photopharmacological targeting of native receptors with genetic precision have recently been reported, providing further opportunities for precise mapping of therapeutic drug action without this limitation.^61–66^

The distinct effects of SNAP-mGluR2 photoactivation in vmPFC-BLA versus pIC-BLA synapses suggest that each projection plays a unique role in defensive responding and avoiding harm. Our data support a role for Grm2^+^ cells in the vmPFC, but not the pIC, in driving spatial avoidance. This central role in avoiding potentially dangerous spatial locations may relate to strong hippocampal afferents to mPFC, which have been previously shown to support spatial avoidance avoidance,^67,68^ whereas pIC and hippocampus are not directly connected.^20^ While our study and others highlight an anxiogenic role of the vmPFC,^15,16^ other studies have reported anxiolytic functions of vmPFC and its outputs,^15,50,51^ and thus it is likely that the vmPFC bidirectionally regulates spatial avoidance. In this study, we demonstrate that diminished neurotransmission in a specific subset of vmPFC-BLA cells reduces anxiety, suggesting an anxiogenic function. Notably, Grm2^+^ projection from the vmPFC were observed only in the BLA, whereas activation of vmPFC projections to the basomedial amygdala (BMA) has previously been demonstrated to reduce spatial avoidance^17^. In accordance, several anatomical studies confirm that the vast majority of vmPFC neurons projecting to BLA reside in upper cortical layers,^52,69–72^ and that BMA receives inputs primarily from vmPFC layer 5 neurons.^17,52^ A lack of Grm2 expression may typify this population of vmPFC-BMA projectors, whereas Grm2 expression marks a subpopulation of exclusively BLA projecting neurons.

Surprisingly, our results also demonstrate a role for Grm2^+^ vmPFC-BLA projections in spatial working memory, as mGluR2 photoactivation in these terminals impaired Y-maze performance. This result suggests a potential circuit basis for the cognitive effects associated with some anxiolytic drugs^73–75^ and the impaired working memory seen with global treatment with mGluR2/3 agonists.^76–78^ This finding highlights the importance of carefully dissecting circuit mechanisms of drug action, as we can begin to ascribe both therapeutic actions and unwanted side effects to specific circuit elements.

Intriguingly, decreased social avoidance and increased exploration were associated with SNAP-mGluR2 photoactivation in pIC-BLA, but not vmPFC-BLA synapses. Both the SIT and FUAT involve sensory stimuli in the form of odors, consistent with the notion that the pIC can assign negative valence to sensory and social cues,^19^ rather than spatial stimuli. The non-significant trend toward anti-social effect seen with Grm2^+^ vmPFC-BLA perturbation is in line with a recent report of a pro-social role for vmPFC-BLA projections,^79^ showing an interesting divergence between a pro-avoidance role for spatial stimuli and anti-avoidance role for social stimuli in this projection. Notably, SNAP-mGluR2 photoactivation in pIC-BLA synapses also decreased avoidance in the NSF task. However, unlike the effects of vmPFC-BLA manipulation which only altered latency to feed in the novel arena, pIC-BLA manipulation was also associated with decreased latency in the home cage. This suggests that the basis of the effect is an increased drive to seek and consume food, consistent with literature showing that IC activity can regulate food consumption^24,80–82^ and the possibility that aberrant insula activity may underlie some eating disorders.^83,84^

Overall, our study points to the pIC-BLA being a potential target for therapeutic intervention for not only treating a subset of anxiety disorders, but also other psychiatric diagnoses like eating disorders, which are defined by abnormal patterns of food consumption, yet for which anxiety is also a dominant clinical symptom.^85^ Our findings are consistent with the emerging picture of the pIC as a general mediator of anxiety-related emotional states^19,23,86^ and suggest that this projection may be particularly promising as a therapeutic target. How might one specifically target this projection in a clinical context? One possibility is to identify novel molecular targets that are specifically expressed or enriched in pIC-BLA. Alternatively, combined brain stimulation and psychopharmacological manipulations may be a viable avenue for leveraging knowledge of circuit-specific pharmacology. Importantly, other projections to the BLA, including those from the aIC^87^ and subcortical regions, likely also contribute to the effects of mGluR2 activation on anxiety-related behavior, motivating future studies.

While distinct recruitments by different sensorial and interoceptive stimuli likely underlie some of the functional differences between Grm2^+^ vmPFC-BLA and pIC-BLA projections, our observations that BLA Grm2^+^ pIC inputs target lateral and vmPFC inputs target medial BLA subregions suggests some circuit-level mechanisms for the distinct behavioral roles of each projection. Several studies indicate that different afferents to the BLA target discrete projecting populations that consequently modulate distinct behaviors.^88–90^ Although transcriptomics studies have only recently started to parse out the molecular complexity of amygdalar neurons,^91–93^ rostro-caudal and medial-lateral subdivisions of BLA have been shown to contain patterns of projection biases.^52,88,90,94–96^ For example, medial BLA neurons project to reciprocally connected neurons in the mPFC,^97,98^ and to the ventral hippocampus. ^52,88,90,95^ Activation of either BLA projections to the mPFC^99^ or ventral hippocampus^100^ is acutely anxiogenic in the OFT and EPM, consistent with the proposed anxiogenic role of Grm2^+^ vmPFC-BLA projections. The medial BLA also sends dense projections to the medial nucleus accumbens (NAc), the bed nucleus of the stria terminalis, and other structures^52^ which may also contribute to the effects of vmPFC-BLA modulation. In contrast, neurons located within lateral portions of BLA do not project to dmPFC nor hippocampus, but extensively target all layers of vmPFC,^52^ ventrolateral areas of NAc, and insular cortex^20^, which may underlie the effects of pIC-BLA perturbations on exploratory, social, and food-seeking behavior. Finally, feedforward inhibition due to parallel recruitment of BLA interneurons modulates synaptic integration in PNs^101,102^ and may further contribute to the differences in the roles of Grm2^+^ vmPFC-BLA and pIC-BLA projections. While we did not explore interneuron diversity^103,104^ in regard to the different cortical inputs in this work, recent studies have highlighted key roles for inhibitory neuron subtypes in information processing and disinhibition in the BLA during various aspects of anxiety and fear. ^102,105–110^

Our data also suggests a role for various forms of synaptic plasticity in underlying the behavioral response to mGluR2 activation. Acute presynaptic inhibition, as seen within seconds in paired pulse measurement in slice recordings, likely contributes to the acute behavioral effects of presynaptic mGluR2 photoactivation *in vivo*. The temporal precision of our approach allowed us to discover that extended activation of presynaptic mGluR2 for twenty minutes *in vivo*, mimicking the *ex vivo* induction of LTD, produced an anxiolytic effect that lasted for at least four hours following receptor de-activation. This points to a long-lasting form of synaptic inhibition but does not directly prove that the same form of LTD seen in slice recordings underlies the behavioral effect. Finally, seven days after mGluR2 photoactivation no behavioral effect was seen, revealing that a homeostatic process, either in the Grm2^+^ vmPFC-BLA synapses or elsewhere in the circuit, re-establishes the prior baseline. The limited desensitization^111^ and protein turnover reported for mGluR2,^112^ suggests that this process is largely driven by downstream intracellular processes. Together these observations point to temporal complexity and feedback regulation in the synaptic effects of cortico-amygdalar GPCR activation. The photopharmacological approach outlined here has the promise to contribute to a deep probing of the relationship between receptor-driven forms of synaptic plasticity and behavioral state changes across many types of receptors, synapses, and neural circuits.

## Methods

### Animals

Grm2-Cre animals were purchased from Mutant Mouse Resource & Research Center (MMRRC) under strain name STOCK Tg(Grm2-cre)MR90Gsat/Mmucd and stock number 034611-UCD, generated from The Gene Expression Nervous System Atlas - GENSAT – Project (NINDS Contracts N01NS02331 & HHSN271200723701C to The Rockefeller University, New York, NY). Wild-type C57BL/6J mice were purchased from Jackson Laboratory. Grm2-Cre animals were bred in house to C57BL/6J mice to create heterozygous Grm2-Cre animals. For all experiments, mice between 8 – 14 weeks of age were used at the start of the experiment. All animal use procedures were performed in accordance with Weill Cornell Medicine Institution Animal Care & Use Committee (IACUC) guidelines under approved protocol (2017-0023).

### AAVs and Chemical Compounds

All adeno-associated viruses (AAVs) were purchased from Addgene or custom-synthesized by the University of Pennsylvania vector core. The following AAVs were used: AAVrg-CAG-FLEX-tdTomato (Addgene 59462-AAVrg), AAVrg-CAG-GFP (Addgene: 37825-AAVrg), AAV5-pCAG-FLEX-EGFP (Addgene: 51502-AAV5), AAV9-EF1a-dfloxed-hChR2-EYFP (Addgene: 20298-AAV9), AAV5-Syn-FLEX-ChrimsonR-tdT (Addgene: 62723-AAV5), AAV9-EF1a-DIO-HA-SNAP-mGluR2-WPRE was previously reported.^45,48^ PORTL compound BGAG_12,400_ was synthesized by J. Broichhagen as previously described. ^48^

### Surgical Procedures

Male and female Grm2-Cre heterozygous mice between 8-14 weeks of age were used unless otherwise noted. The following coordinates from skull were used for AAV or BGAG_12,400_ injections (AP, ML, DV): BLA (-1.7, ±3.3, -4.65), vmPFC (1.95, ±0.35, -2.65) and pIC (-0.50, ±4.05, -4.05). For all surgical procedures mice were anesthetized using 1.5-2% isoflurane and were injected using a Kopf stereotaxic and World Precision Instruments (WPI) microinjection syringe pump with a 10 mL syringe and 33g blunt needle at 50-100 nL/min, with the syringe left in place for a further 5-10 minutes to allow diffusion.

For retrograde neuroanatomy, 250 nL each of AAVrg-CAG-FLEX-tdTomato and AAVrg-CAG-GFP (500 nL total pre-mixed) was injected unilaterally into BLA and expressed for 5-6 weeks. For slice electrophysiology and *in vivo* optogenetics, 250 nL of AAV5-EF1a-doublefloxed-hChR2(H134R)-EYFP-WPRE-HGHpA and/or AAV5-Syn-FLEX-rc[ChrimsonR-tdTomato] were injected bilaterally into the vmPFC or pIC and expressed for 5-7 weeks. For *in vivo* mGluR2 photoactivation, 500-600 nL of AAV9-EF1a-DIO-HA-SNAP-mGluR2-WPRE was co-injected with a 1:20 dilution of AAV5-pCAG-FLEX-EGFP-WPRE expressed for 5-7 weeks.

For intra-BLA cannulation, ICV Single Dummy Cannulas (26 Gauge Stainless-Steel) from P1 Technologies Inc. at 0.5 mm length and 0.1 mm thickness were bilaterally implanted to target BLA at AP -1.75 and ML +/- 0.35 from bregma. Cannulas were adhered to the skull using C & B Metabond (Parkell), and experiments were conducted 2 weeks later.

For behavioral photopharmacology and optogenetics, dual fiber optic/cannula implants manufactured by Doric Lenses were used as previously described^45^ but specifically designed to target BLA (AP -1.7, ML +/- 3.3, DV -4.5) and implanted 2 weeks before experimental testing began. Implants were adhered to the skull using C & B Metabond (Parkell). 14-20 hours before each behavioral test, mice were anesthetized using 1.5-2% isoflurane and dual opto-cannula guide plugs were replaced with 100 mm inner diameter micro-injectors to target BLA (AP -1.7, ML +/- 3.3, DV -4.6). 1 µL per hemisphere of 25 µM BGAG_12,400_ was infused through micro-injectors (outer diameter = 166 mm, inner diameter = 100 mm) at 100 nL per minute using polyethylene tubing connected to a Harvard Apparatus (Harvard Biosciences, Inc.) 11 Elite dual syringe infusion pump and 10 mL Hamilton Company. After infusions, micro-injectors were replaced with optical fibers (outer diameter = 200 µm, numerical aperture = 0.66) to target BLA (AP -1.7, ML +/- 3.3, DV -4.55). All AAV injections and implants were validated post-hoc via histology and mice are excluded from analysis if they deviated from the defined target coordinates.

### Brain Slice Electrophysiology

Mice were deeply anesthetized with isoflurane and perfused with ice-cold NMDG-HEPES aCSF (in mM): 93 NMDG, 2.5 KCl, 1.2 NaH_2_PO4, 30 NaHCO_3_, 20 HEPES, 25 Glucose, 5 sodium ascorbate, 2 thiourea, 3 sodium pyruvate, 10 MgSO_4_, 0.5 CaCl_2_. BLA coronal slices (300 μm) were collected in the same solution and kept for 30 min at 34 °C, followed by at least 45 minutes at room temperature in a modified HEPES aCSF containing (in mM): 92 NaCl, 2.5 KCl, 1.2 NaH_2_PO_4_, 30 NaHCO_3_, 20 HEPES, 25 Glucose, 5 sodium ascorbate, 2 thiourea, 3 sodium pyruvate, 2 MgSO_4_, 2 CaCl_2_. Solutions were pH-corrected to 7.4, 300-305 mOsm and were continuously bubbled with 95% O_2_/5% CO_2_. Recordings were performed at 29-31° C in a standard oxygenated aCSF containing (in mM): 124 NaCl, 2.5 KCl, 1.2 NaH_2_PO_4_, 24 NaHCO_3_, 5 HEPES, 12.5 Glucose, 1.3 MgSO_4_, 2.5 CaCl_2_. Whole-cell patch-clamp recordings were performed with pipettes of 3-7 MΩ resistance, filled with an intracellular solution containing (in mM): 120 CsMeSO_4_, 8 NaCl, 0.3 EGTA, 10 Tetraethylammonium-Cl, 10 HEPES, 2 Mg_2_-ATP, 0.3 Na_2_-GTP, 3 QX-314-Cl and 13 biocytin (pH 7.3 and 285–290 mOsmol / kg). Recordings were performed in voltage-clamp at -80 mV on a computer-controlled amplifier (MultiClamp 700B Axon Instruments, Foster City, CA) and acquired with an Axoscope 1550B (Axon Instruments) at a sampling rate of 10 kHz and low-pass filtered at 1 kHz. In some experiments tetrodotoxin (TTX-Tocris, #1078) was applied at 1 µM for monosynaptic mapping experiments and LY379268 (LY37 – Tocris #2453) was applied at 100 nM for mGlur2/3 activation.

Cells were classified as pyramidal neurons (PNs) according to their triangular somatic shape with all other cells were considered interneurons (INs). Pilot experiments were performed in current clamp and neuronal firing properties could be recorded and confirmed to match expected cell classes. All cells were filled with biocytin for post-hoc confirmation of morphology and location within BLA. Data from two neurons were not included due to undistinguishable identity. Data were analyzed in Clampfit, Microsoft Excel, and Prism (Graphpad Software, Inc.).

A CoolLED pE-400 coupled to the microscope was used for visualization and photo-stimulation with light intensities of 1-2 mW/mm^2^. Under voltage-clamp mode, oEPSC were measure after opsin stimulation via 1-2 ms 470 nm (ChR2) or 590 nm (ChrimsonR) pulses. To ensure assessment of monosynaptic inputs, latency was used as a first checking point of analysis: onsets above 4.1 ms were not considered for further analysis^101,113^. Amplitude (pA) of oEPSC and PPR was measure from an average of 5-10 sweeps. For LTD, opsin stimulation every 30s was maintained and a stable baseline for ≥10 min was established prior to LY37 application. For dual input experiments (**Fig. 6**), 590 nm stimulation did not detectably activate ChR2 but 470 nm pulses occasionally weakly activated ChrimsonR to produce oEPSCs of ∼10-15 % of the response to ChR2^113^.

### Behavioral Testing

All mice were acclimated to the behavioral room for 1 hr before each behavioral test and handled for 3-5 consecutive days before their first behavioral test. All apparatuses were cleaned with 70% ethanol between each animal unless except for FUAT, where arenas were cleaned with Clorox Wipes.

For pharmacology experiments, 8-12 week old WT C57BL/6J mice or WT Grm2-Cre-(Grm2-Cre+ littermates) were used. Mice were intraperitoneally injected with 3 mg/kg (diluted in saline) LY34 or LY37 or saline 30 min before the behavioral test. For local infusion experiments, mice were infused with 100 nL of 10 mM LY37 diluted in saline or saline into the BLA at 100 nL/min 5 min before behavioral testing. LY37 or saline was delivered via implanted cannula using polyethylene tubing connected to a Harvard Apparatus (Harvard Biosciences, Inc.) 11 Elite dual syringe infusion pump and 10 mL Hamilton Company.

For mGluR2 photoactivation experiments, male Grm2-Cre^+^ mice were tested starting 7 weeks after viral injection. Light was delivered via LED as previously described. ^45,48^ Briefly, dual fiberoptic cannulas were connected to a dual fiberoptic patch cord (200 μm diameter per cord, Doric) to a fiberoptic rotary joint (Doric) and then attached to a mono fiberoptic patch chord (400 μm, Doric) connected to a 4 Channel LED Driver (LEDD_4) and a 12 V Fan Power Adaptor (FPA). The LED unit was manipulated via Doric Neuroscience studio software (DORIC STUDIO V5.3.3.6). For ON photoswitching, ∼10 mW/mm^2^ 385 nm light was pulses for 2 s ON followed by 2 s OFF 3 times every 48 s and repeated continuously until the behavior was completed. *In vivo* light stimulation using the 385 nm stim protocol is not expected to exceed a firing temperature change threshold of 1 degree C based on Stujenske et al^114^. ON photoswitching commenced 5 min before the start of the behavioral test either during the habituation phase or in the home cage of the animal, except for OFT where the light was started 5 min after the test began. For *in vivo* behavioral plasticity experiments (**Fig. 4**), the light protocol started from the beginning to the end of the 20 min OFT-stimulation session. No light animals received the same treatment except the LED was not turned on. For behavioral battery, mice underwent tests one week apart in the following order: OFT, EPM, SIT, NSF, FUAT, YMT, TST (see **Fig. 5**). After the behavior was complete, mice received a 5 second pulse of 15 nm light at ∼10 mW/mm^2^ to relax BGAG_12,400_ to its inactive *trans* form. For channelrhodopsin EPM experiments (**Fig. 4**), blue light (465 nm) was delivered at 10Hz, with 5 ms pulses and mice were tested in EPM for a period of 15 minutes divided into ‘light off’(5min), followed by ‘light on’(5min) and again ‘light off’ (5 min) epochs.

#### Open Field Test (OFT)

Mice were placed in the corner of a 50 cm x 50 cm open field arena with a grey Plexiglas flooring and black Plexiglas walls 35 cm high, with a bright light above with a LUX of 300 and a camera overhead for recording. Mice freely explore the arena for 20 minutes, and time spent and entries into the center of the arena is quantified (center = middle 25% of the arena). Plotted data corresponded to the last 5 min of the test. Activity was recorded and analyzed using SMART and Ethovision analysis software.

#### Elevated Plus Maze (EPM)

Mice were placed in the closed arm of the elevated plus arena constructed of grey Plexiglas, which was raised 70 cm above the floor and consisted of two opposite closed arms with 14-cm-high opaque walls and two opposite open arms of the same size (29 cm × 6 cm). A bright light was placed directly above (250 LUX) the center of the maze with a camera overhead for recording behavior. Mice were tested for 5 min and time, entries, dwell time, and head dips in the open arms were recorded post-hoc using Ethovision and blind hand scoring. For a cohort performed for c-fos expression quantification in BLA (Fig. S4), mice were perfused 90 min following EPM and light stimulation.

#### Social Interaction Test (SIT)

A three compartment apparatus consisting of a white Plexiglas rectangular box (57.15 cm x 22.5 cm x 30.5 cm), without a top was used. The 3 compartments are of equal size, with an insertable door between the middle chamber and the 2 end chambers. Plexiglas cups with bars 1 cm apart around were placed in each side of the end compartments during testing sessions. During the habituation phase, mice were freely allowed to explore all 3 chambers with empty cups for 5 minutes. Mice were then moved back into the middle chamber and doors were inserted to keep the mouse there. The cups were then filled with a novel object and an age and sex matched littermate mouse that was novel to the testing phase with a partner mouse but had previously been habituated to the cup (side of arena randomized per trial and group). The chamber doors were then removed, and mice were allowed to freely explore all 3 chambers while their activity was recorded. Time spent in social zone, which was the half of the social compartment where the cup with the social mouse was, was calculated post-hoc using Ethovision. Social interaction ratio was calculated as (time in social zone)/(time in social zone + time in object zone). Object zone is defined the same as the social zone but in the object side of the chamber.

#### Novelty Suppressed Feeding (NSF)

Mice were food deprived for 16 hr overnight and then placed in the corner of a 50 cm x 35 cm white plastic container with wood chip bedding on the bottom. In the center of the arena, a large (∼5 g) food pellet was placed. Direct bright light and a camera were fixed overhead (LUX ∼500). Latency to eat the food pellet was recorded for up to 5 min, where once the mouse eats the food pellet they are removed from the arena. After the arena latency test, mice were returned to their (empty) home cage with another large food pellet placed in the center. Latency to eat the food pellet in the home cage was recorded for 5 min. Data was recorded in real time using a stopwatch, distance traveled was analyzed post-hoc using Ethovision.

#### Fox Urine Avoidance Task (FUAT)

Mice were placed in the corner of an empty home cage inside a hood with light overhead (LUX ∼300), and a white filter paper with 10 μL of water was placed in the opposite corner. Activity was recorded for 5 min, including time in the water zone (cage divided into 3 equal zones) and sniffing time. After a 5 min habituation, the water filter was replaced with a filter paper with 10 μL of 1% TMT (Sigma, CAS:13623-11-50). Mice were then recorded for an additional 5 min to analyze the test trial with TMT. Time in each zone was calculated using Ethovision and hand scoring.

#### Y-Maze Test (YMT)

Mice were transferred to a Y-maze consisting of three equally sized plastic arms for a 5-minute test as previously described. ^45,48^ Lights were turned on at full power in the room (LUX ∼300) and a camera was placed overhead to record. Correct and incorrect alternations (total alternations) were recorded by the experimenter in real time.

#### Tail Suspension Test (TST)

Mice were hung by the tail on a metal bar inside a 45 x 75 cm black plastic box with walls subdividing each animal into 45 x 15 cm individual cubes for 6 min. Time immobile was calculated using SMART real time analysis, which was defined as less than 100 cm^2^/second using the automated software.

### Histology and Microscopy

After *in vivo* experimental procedures, mice underwent transcardiac perfusion with PBS, followed by 4% cold paraformaldehyde. Brains were extracted and kept in 4% paraformaldehyde for 24 hours followed by PBS solution until sectioned. Brains used for c-Fos staining were further transferred to 30% sucrose overnight, then embedded in optimal cutting temperature compound (OCT, Histolab Products AB or Tissue-tek) and stored at -80°C until cryostat sectioned (40 µm sections). Brains were mounted on a Leica Vibratome in cold PBS solution and sliced at 60 µm for immunohistochemistry (IHC) and 70 µm for all other histology analysis. Slices were wet mounted to glass slides and secured with coverslip using VECTASHIELD HardSet Antifade Mounting Medium with DAPI (Vector Laboratories). Glass slides were imaged using an Olympus Confocal FV3000 or an LSM-880 confocal microscope and images were processed using ImageJ.

We performed IHC for HA (HA-SNAP-mGluR2) and c-Fos in coronal sections. Sections were placed in boiling antigen retrieval solution (10 mM Citric Acid 0.05% Tween-20, pH 6) for 30 min. They were washed 2 x 10 min in PBS, permeabilized 3 x 10 min in PBS/0.1% Tween20 and blocked for 1 hr in dilution buffer (50 ml PBS, NaCl 0.5 M, BSA 2.5%, Tween20 0.3%) with 5% normal donkey serum (Labome, 017-000-121) at room temperature. Sections were incubated at 4° C overnight in 1:500 recombinant rabbit anti-HA tag (Abcam, AB236632) or rabbit anti-c-Fos (Abcam, AB190289) antibody diluted in dilution buffer. The next day, sections were washed 4 x 15 min with PBS/0.1% Tween-20, incubated 1 hour at room temperature in 1:500 Alexa Fluor-conjugated antibody (Invitrogen, Alexa Fluor 568 #A10042, or Alexa Fluor 488 #A-21206) in dilution buffer, and washed 3 x 15 min with PBS/0.1% Tween20. Sections were incubated for 10 min at room temperature in 1:5000 DAPI (Thermo Fisher, D1306) in PBS, mounted (Vector Laboratories, H-1400-10 or Fluoromount-G, SouthernBiotech), and stored at 4° C until imaging.

### Quantification and Statistical Analysis

Retrograde neuroanatomy experiments included data from at least 3 separate mice. Triplicate images of each brain region were taken, and results across the 3 brain regions were averaged to determine number of cells per animal for eGFP, tdT, and DAPI channels. Brain region boundaries were cropped from images in ImageJ using a reference brain atlas. For counting, brain regions were cropped based on Atlas coordinates and morphology for specific subregions. Images were split by channel for DAPI, eGFP, and tdT. DAPI was counted via ImageJ automated analysis, where images were first background subtracted (∼200) and then normalized to the Threshold analysis. Then, images were put through the “Fill Holes” and “Watershed” functions, and finally particles were analyzed with Analyze Particles function with parameters size 0-50 microns^2^. GFP and tdT channel overlap was manual counted using cell counter. After individual cell counting and ImageJ automated DAPI counting and brain region size analysis, quantification of parameters was completed in Excel and visualized using Graphpad.

Behavioral experiments were analyzed with SMART (Harvard Apparatus), Ethovision (Noldus) or blinded hand scoring as indicated. Mice were excluded from tests based on outlier calculation using Graphpad Prism. Avoidance Z-scores for each experimental cohort were calculated by normalizing each +Light (385 nm) group to its respective control (no-Light) group. Z-score of each mouse corresponded to = (Mouse Score – Control Average)/Standard Deviation of Control Group. Parameters used for each behavior were: % Time in Center of bin 15-20 min (OFT), % Time Open Arms (EPM), % Time Social Chamber (SIT), Latency to feed (NSF), Ratio Fox Urine/Water exploration time (FUAT).

Radar plots (**Fig. 5D**) were generated in Microsoft Excel using Z score calculation above for each test, with locomotion calculated using an average of the locomotion data between OFT, EPM, SIT, NSF, FUAT, and YMT. Tissue heating plots were generated using Matlab custom scripts and optical fiber and LED parameters as stated above.

Statistical analysis was performed using Microsoft Excel and Prism (GraphPad Software Inc.) for all experiments. Individual statistical methods are indicated in the figure legends. Statistical significance is indicated in the text and figure legends as ∗p < 0.05, ∗∗p < 0.01 and ∗∗∗p < 0.001. No additional methods beyond the statistical tests stated in the text and figured legends were used to determine whether the data met assumptions of the statistical approach.

## Acknowledgements

We are grateful to Francis Lee (Weill Cornell Medicine) and Kristen Pleil (Weill Cornell Medicine) for access to behavioral equipment and valuable discussions and to Prerana Vaddi for technical assistance. This work is supported by NIH grants R01MH129693 (J.L.), R01MH123154 (C.L.), R01MH118451 (C.L.), R01NS126073 (C.L.), F31MH123130 (V.G.), K08MH127383 (J.M.S.), the Swedish Research Council VR2020-06395 (H.M.), the Brain & Behavior Research Foundation - NARSAD Young Investigator Grant BBRF-3130 (H.M.), funding from the European Union’s Horizon Europe Framework Programme (deuterON, grant agreement no. 101042046 to J.B.), the Rohr Family Research Scholar Award (J.L.), the Monique Weill-Caulier Award (J.L.), the Hope for Depression Research Foundation (C.L.), and the Pritzker Neuropsychiatric Disorders Research Consortium (C.L.).

---

**Figure S1.**
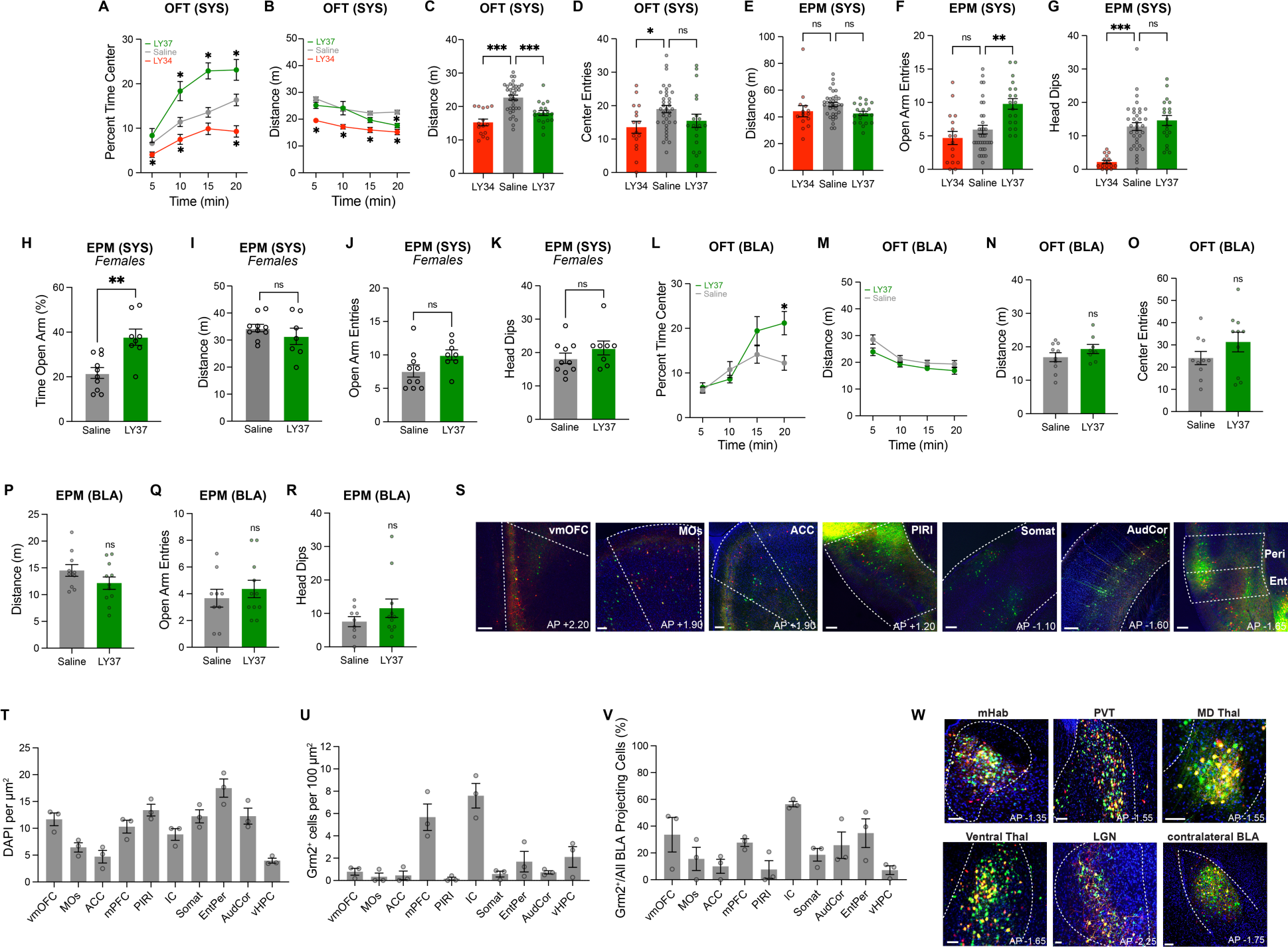
Further analysis of behavioral effects of mGluR2/3 agonism and mapping of Grm2^+^ amygdalar projections. **(A-G)** Effects of I.P. injection of LY37 or LY34 on spatial avoidance in the OFT (A-D) and EPM (E-G). **(H-K)** I.P. injection of LY37 produces a similar anxiolytic effect in female mice as seen with male mice (see Fig. 1) in the EPM. **(L-R)** Effects of intra-BLA infusion of LY37 on spatial avoidance in the OFT (L-O) and EPM (P-R). **(S)** Representative images showing BLA-projecting cells across cortical regions. **(T-V)** Summary bar graphs showing density of all cells (T), density of Grm2^+^ BLA-projecting cells (U), and proportion of BLA projections cells that are Grm2^+^ (V) across cortical regions. **(W)** Representative images showing BLA-projecting cells across subcortical regions. Points represent individual mice. For (A), (B), (L), 2-way ANOVA was used; for (C-G), 1-way ANOVA was used; for (H-K), (N-R) unpaired t-test was used. All data shown as mean ± SEM; * P<0.05, ** P < 0.01, *** P < 0.001. Scale bars are 200 µm.

**Figure S2.**
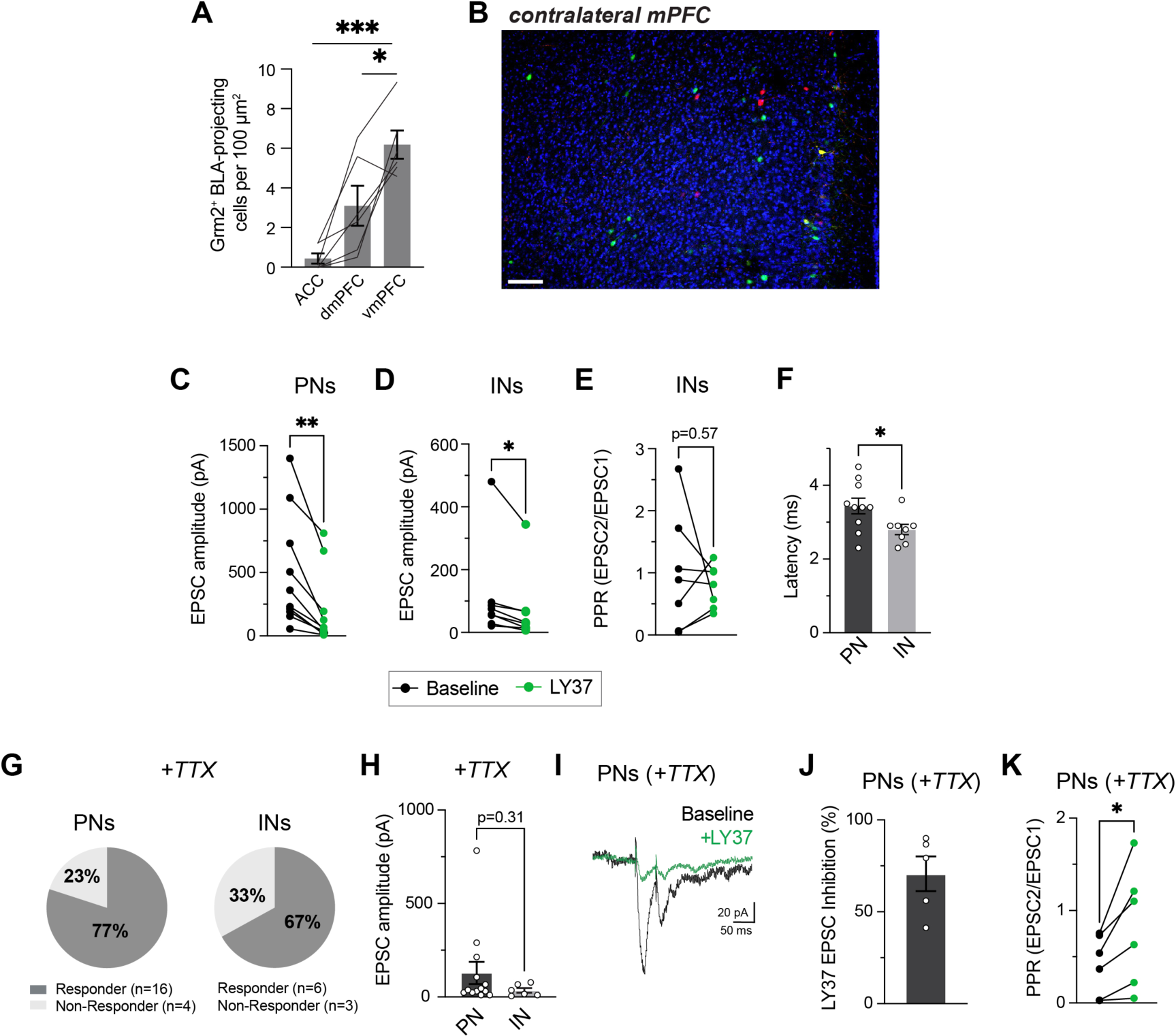
Further analysis of Grm2^+^ projections from PFC to BLA. **(A)** Summary bar graph showing density of Grm2^+^ BLA-projecting cells across the mPFC. **(B)** Representative image showing very sparse population of BLA-projecting cells in contralateral mPFC. **(C-D)** Quantification of effects of LY37 application on oEPSCs from Grm2^+^ vmPFC projections onto PNs (C) and INs (D). **(E)** Quantification of effects of LY37 application on paired pulse ratio of oEPSCs onto INs. **(F)** Summary graph showing latency from optical stimulation to oEPSC onset in PNs versus INs. **(G-K)** In the presence of TTX Grm2^+^ vmPFC-BLA connectivity is seen with most PNs and INs (G, H) and oEPSCs onto PNs are highly sensitive to LY37 (I) in terms of both amplitude (J) and paired-pulse ratio (K). Lines represent individual mice (A) and points represent individual cells. For (A) 1-way ANOVA was used; for (C-F), (K) paired t-test was used; for (F), (H) unpaired t-test was used. All data shown as mean ± SEM; * P<0.05, ** P < 0.01, *** P < 0.001. Scale bars are 200 µm.

**Figure S3.**
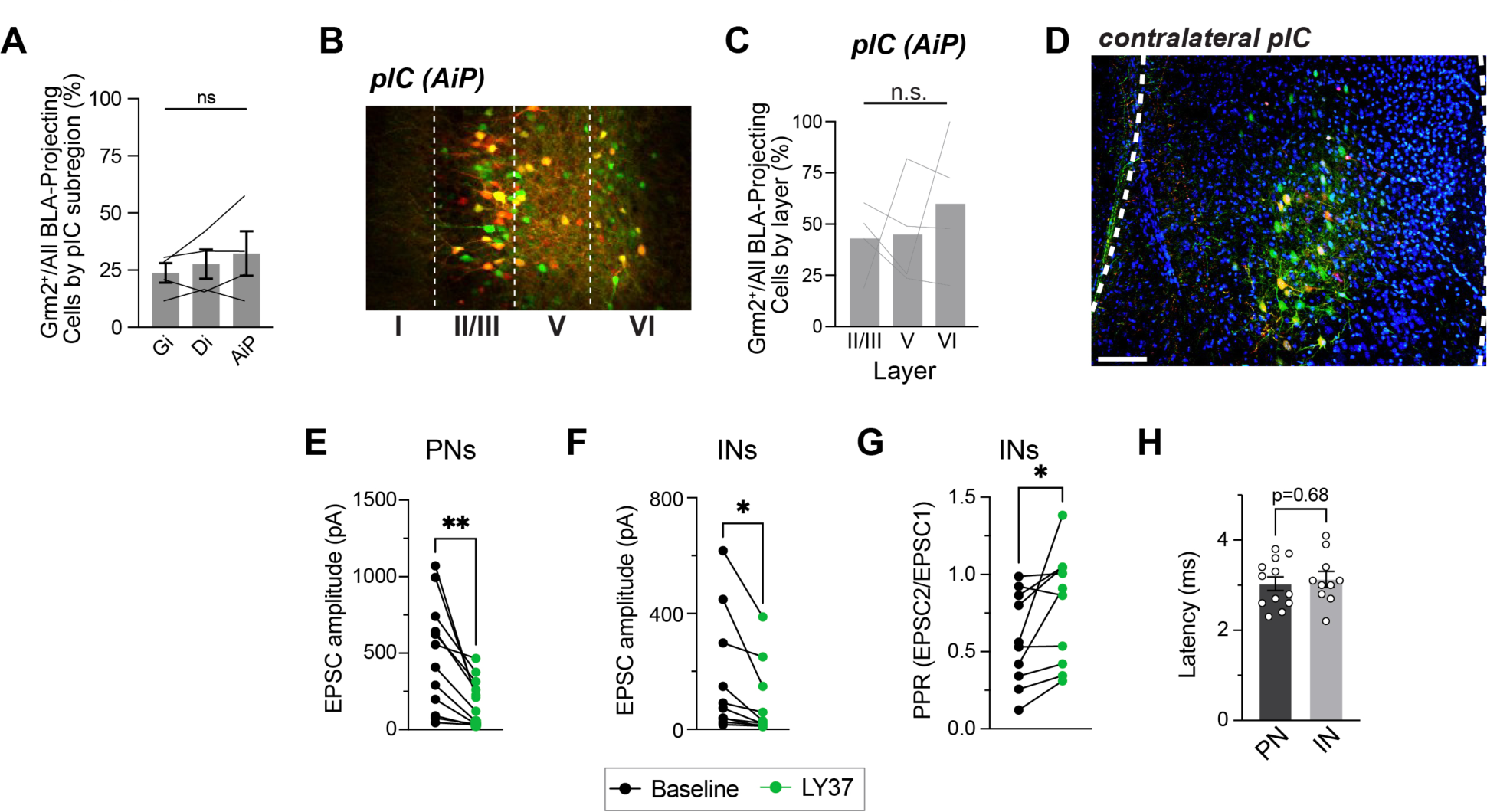
Further analysis of Grm2^+^ projections from IC to BLA. **(A)** Summary bar graph showing proportion of BLA-projecting cells that are Grm2^+^ across pIC subregions. **(B-C)** Representative image and summary graph showing that Grm2^+^ BLA-projecting cells show a similar layer distribution to the total population of BLA-projecting cells. **(D)** Representative image showing substantial population of BLA-projecting cells in contralateral pIC. **(E-F)** Quantification of effects of LY37 application on oEPSCs from Grm2^+^ pIC projections onto PNs (E) and INs (F). **(G)** Quantification of effects of LY37 application on paired pulse ratio of oEPSCs onto INs. **(H)** Summary graph showing latency from optical stimulation to oEPSC onset in PNs versus INs. Lines represent individual mice (A), (C) and points represent individual cells. For (A), (C) 1-way ANOVA was used; for (E-G) paired t-test was used; for (H) unpaired t-test was used. All data shown as mean ± SEM; * P<0.05, ** P < 0.01. Scale bars are 200 µm.

**Figure S4.**
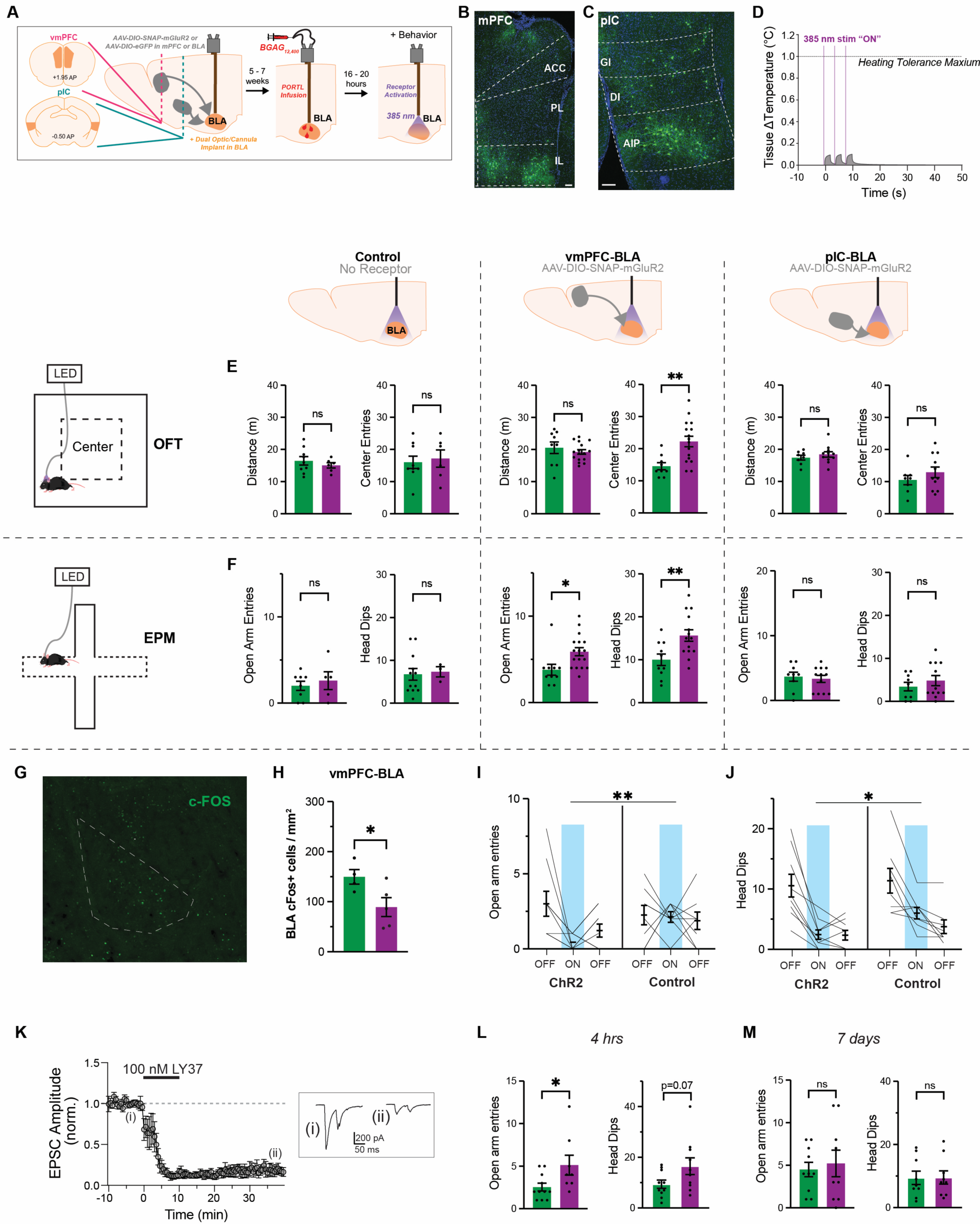
Further behavioral and functional analysis of Grm2^+^ cortico-amygdalar projections. **(A)** Schematic showing experimental details and timeline for projection-targeted behavioral photopharmacology. **(B-C)** Representative immunohistochemistry images showing SNAP-mGluR2 expression in vmPFC (B) and pIC (C) in Grm2-Cre mice. **(D)** Graph of simulation of tissue heating *in vivo* using the 385 nm stim protocol, predicting that the BLA does not exceed a firing temperature change threshold of 1 degree C. **(E-F)** Further analysis of the effects of SNAP-mGluR2 photo-activation on avoidance behavior in the OFT (E) and EPM (F). **(G-H)** c-Fos staining of BLA in mice expressing SNAP-mGluR2 in vmPFC with (magenta) or without (green) 385 nm stimulation during EPM reveals a decrease in positive cells due to mGluR2 activation. **(I-J)** Further analysis of behavioral effects of ChR2 stimulation in Grm2^+^ vmPFC-BLA terminals in the EPM. **(K)** Optogenetics-assisted slice electrophysiology showing that LY37 produces long-term depression in Grm2^+^ vmPFC-BLA projections. Inset (bottom) shows representative traces before and after 10 min treatment with LY37. N=6 cells from 3 mice. **(L-M)** Further analysis of effects of activation of presynaptic SNAP-mGluR2 in Grm2^+^ vmPFC-BLA terminals four hours (L) or 7 days (M) prior to EPM. Points and lines represent individual mice. OFT = open field test, EPM = elevated plus maze. For (E), (F), (H), (L), (M) unpaired t-test was used; For (I), (J) 2-way ANOVA was used. All data shown as mean ± SEM; * P<0.05, **P<0.01,. Scale bars are 20 µm.

**Figure S5.**
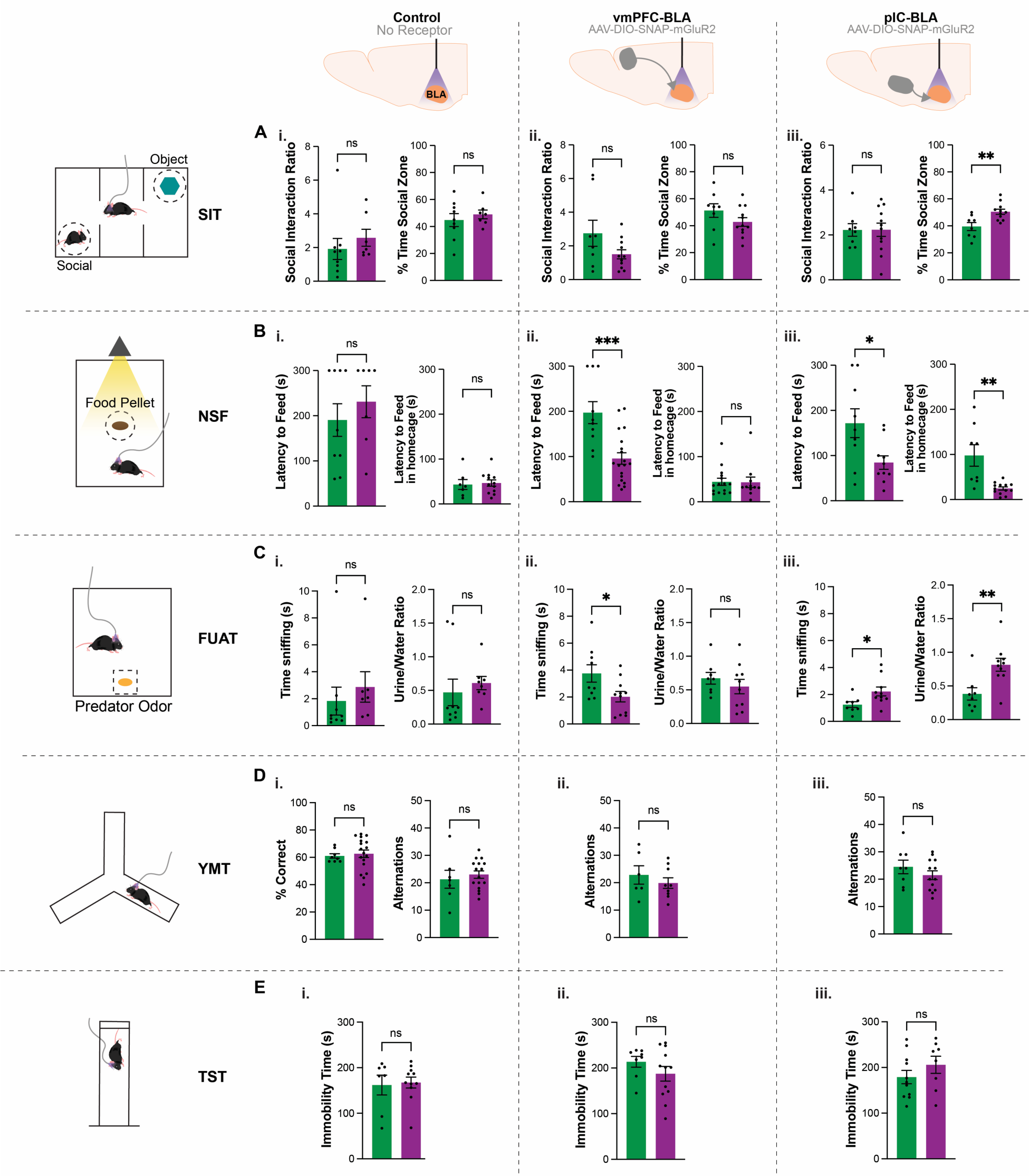
Further behavioral analysis of Grm2^+^ cortico-amygdalar projections across avoidance behaviors. Further analysis of the effects of SNAP-mGluR2 photo-activation on avoidance behaviors across assays. Points represent individual mice. SIT = social interaction test, NSF = novelty suppressed feeding, FUAT = fox urine avoidance test, YMT = Y Maze Test, TST = tail suspension test. Unpaired t-test was used. All data shown as mean ± SEM; * P<0.05, **P<0.01, ***P<0.001.

**Figure S6.**
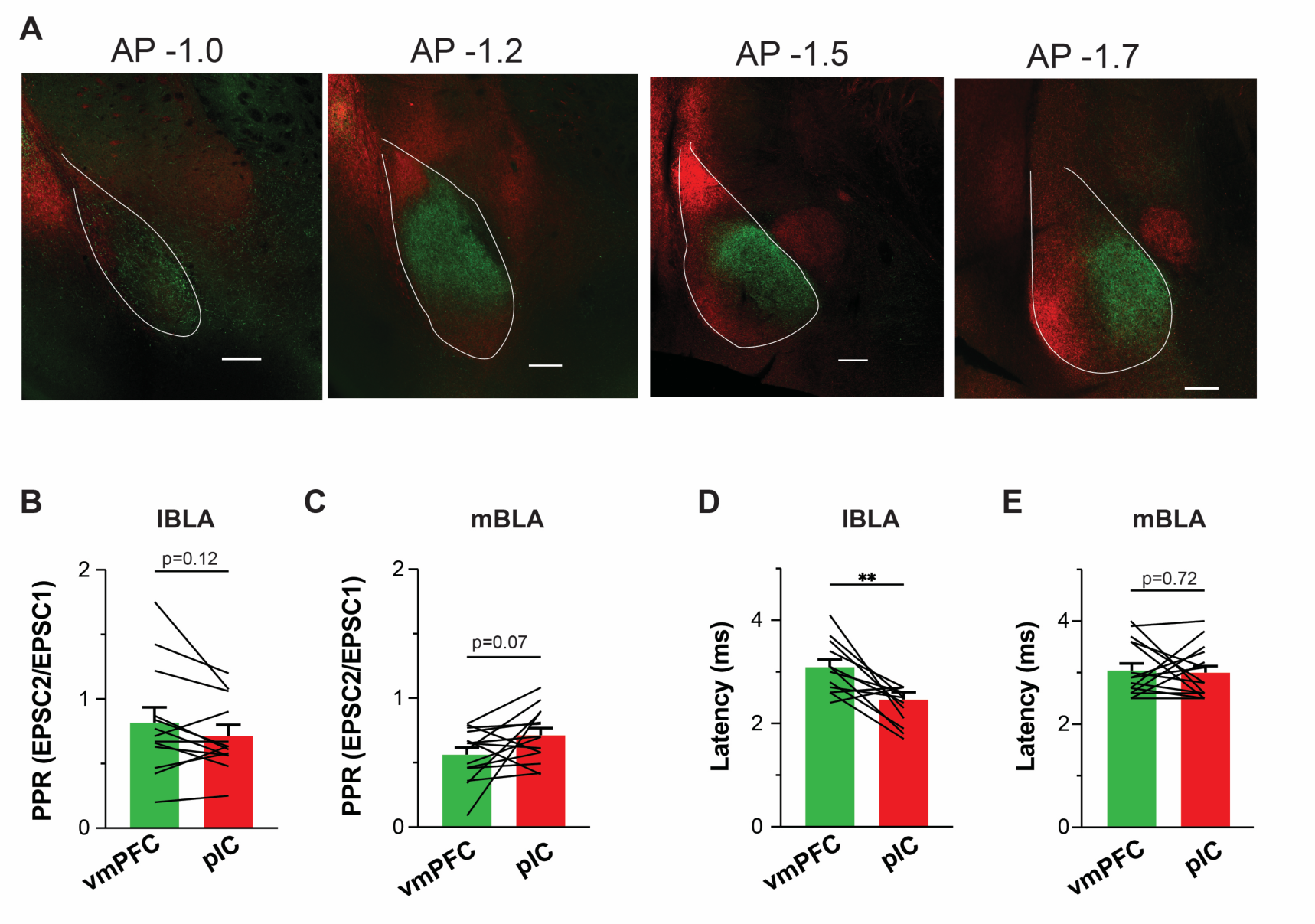
Further comparative anatomical and functional analysis of Grm2^+^ cortico-amygdalar projections. **(A)** Representative images showing spatial segregation of Grm2^+^ vmPFC (green, YFP-ChR2) and pIC (red, tdT-Chrimson) inputs to the BLA. **(B-C)** Summary bar graph showing paired pulse ratios for oEPSCs from both cortical inputs for lateral (B) and medial (C) BLA. **(D-E)** Summary graph showing latency from optical stimulation to oEPSC onset in PNs from both cortical inputs for lateral (D) and medial (E) BLA. Lines represent individual cells. Paired t-test was used. All data shown as mean ± SEM; Scale bars are 200 µm.

